# Defective HIV DNA genomes provide ancestral relevance critical for phylogenetic inference of reservoir dynamics

**DOI:** 10.1101/2022.05.04.490630

**Authors:** Lauren E Droske, Andrea S Ramirez-Mata, Melanie N Cash, Jose Estrada, Stephen D Shank, Adam Browning, Faezeh Rafiei, Sergei L Kosakovsky Pond, Marco Salemi, Brittany Rife Magalis

## Abstract

During the course of infection, human immunodeficiency virus (HIV) maintains a stably integrated reservoir of replication-competent viruses within the host genome that are unaffected by antiretroviral therapy. Curative advancements rely heavily on targeting the anatomical reservoirs, though determinants of their evolutionary origins through phyloanatomic inference remain ill-supported through current sequencing and sequence analysis strategies. The vast replication-defective genomic landscape that comprises the HIV DNA population is often discarded in these evolutionary endeavors, despite key information regarding competent ancestry that can be gained from captured genomic regions outside the historically used viral envelope gene. Here, we describe the application of small-amplicon, single-cell DNA sequencing to blood and lymph node samples from a treatment-interrupted S[imian]IV-infected animal model and evaluate the contribution of genome coverage and inclusion on phylogenetic resolution and phyloanatomic inference. Findings from this study point to incomplete genomes as a significant source of phylogenetic information on movement of virus between tissue reservoirs during therapy.

## Introduction

Though the advent of combined antiretroviral therapy (cART) has vastly improved survival rates of persons living with HIV (PLWH), complete clearance of the virus has yet to be attained^1,2^. Incomplete viral eradication is in large part due to the ability of HIV, as with other retroviruses, to maintain a form of latency through integration of the viral genome into the cellular genomic deoxy-ribonucleic acid (gDNA)^3–5^. With the exception of rare elite controllers of viremia, viral rebound to levels approximating initial infection can result from the interruption of cART, indicating the production of new virions from the drug-impervious viral reservoir^6^. More recent investigation of the HIV+ DNA population within the infected host has demonstrated that the vast majority (*≥* 93%) actually consists of incomplete, replication-incompetent virus generated through a variety of steps in the viral replication process^7–11^, posing a needle-in-a-haystack issue for pinpointing reservoir origins of viral rebound.

Statistical phylogenetic inference of movement of pathogen populations between distinct locations has been applied at both a global (phylogeography^12^) and anatomical scale (phyloanatomy^13–15^). These approaches are based on the principle of shared common ancestry between samples taken at varying time points and locations in the form of a phylogenetic tree, or phylogeny^13–15^. Given known states (i.e., tissue sampled) supplied for the leaves of a phylogeny, statistical reconstruction of ancestral states at the internal nodes can be carried out, allowing for inference of the frequency of “transitions” between states along the branches of the tree, often interpreted as migration. HIV phyloanatomic studies have primarily relied on sequence information derived from the viral envelope (*env*) gene region, specifically that encoding the surface glycoprotein (*gp120*), owing to its abundant evolutionary signal necessary for reliable phylogenetic reconstruction^14,16,17^. Reconstruction of discrete sampling states within the HIV *env* phylogeny from infected tissue donors as part of the Last Gift cohort, Chaillon *et al.* (2020)^18^ described specific deep-tissue reservoirs and viral migration pathways following cessation of therapy. However, restriction of phylogenetic analysis to a single gene or intact virus may result in loss of mutational signal potentiating erroneous conclusions regarding both reconstruction of the phylogeny and ancestral states within the phylogeny. In other words, though replication-defective, incomplete genomes are the progeny of replication-competent viral ancestry and thus share mutational history that are represented in the phylogeny and may be critical missing pieces in the migration history. The improved sequence resolution potentially provided by the utilization of all sampled viral genomes, regardless of their replication competency, may act to increase reliability and interpretability of phylogenetic inference, fundamental to elucidating the origins of the HIV reservoir and viral rebound in space and time^19–21^.

Droplet-based microfluidic approaches to single-cell isolation are well-suited for reliable whole-genome sequencing (WGS) of heterogeneous populations (e.g., Eastburn *et al.*^22,23^) and have been instrumental in tumor cell population characterization through targeted host DNA sequencing, beginning with Pellegrino *et al.* (2018)^24^. This microfluidic approach relies on picoliter-volume droplets of aqueous biological reagents encapsulated in an oil-based emulsion^25^, enabling massively parallel polymerase chain reaction (PCR) with ultra high-throughput sequencing. Although attempts have been made to improve HIV reservoir characterization through alternative WGS techniques, they fall short of reliable completeness and/or depth, can be labor-intensive, and/or introduce several *a priori* assumptions^26–28^, such as inclusion of critical sites among all sequences^9,29^. Most notably, Sun *et al.*applied single-cell DNA sequencing (scDNA-seq) to HIV-infected cell populations, utilizing sorting of short-range PCR-positive amplification of two genomic regions, enriching for proviruses with the potential for infectivity^30^. Sun *et al.* expanded on this protocol to detect an increased number of defective viral genomes in 2023^31^, though still only including 18 targeted regions (200 − 300 base pairs in length) across the HIV-1 genome. Moreover, both studies enriched for CD4+ T cells. While resting CD4+ T lymphocytes represent the largest and most well-characterized harbor for intact virus, there is mounting evidence from both animal models and human donors that other cell types contribute to the total HIV reservoir^26^. Lastly, these approaches have not been used to distinguish episomal DNA from integrated DNA, which may act as a surrogate for active or recent infection^32,33^ (though still debated^34^). Upon integration disruption, whether through antiretrovirals or stochastic processes, the viral DNA can circularize via ligation of the long terminal repeat (LTR) regions flanking the genome. This process results in LTR sequence junctions that can be detected using targeted sequencing to better understand their contribution to ongoing viral evolution. The studies by Sun *et al.*, as well as others^35^, have largely focused on characterization of the expansion of the reservoir through cellular replication; however, the utility of this technique in phylogenetic and phyloanatomic inference has not been determined.

Rhesus macaques infected with pathogenic simian immunodeficiency virus (SIV) offer an ideal animal model for controlled sampling of otherwise invasive tissues in human beings for rigorous identification of tissues and cell types that contribute to ongoing viral evolution in the face of therapy and viral rebound following therapy interruption^36^. In this study, peripheral blood mononuclear cell (PBMC) and lymph node (LN) samples were taken from three animals (HG02, JA41, and JP70) either during therapy (50-120 days post-inoculation [dpi]) or post-therapy interruption at necropsy (197 dpi) (Supplementary Figure S1). We applied non-enriched scDNA-seq to these samples, targeting 37 regions across the SIV genome to determine the level of signal contributed by low-coverage genomes in the inference of viral reservoir dynamics across the blood and lymph node compartments.

The scDNA-seq approach described herein couples droplet microfluidic technology and next-generation sequencing to capture near-full-length (NFL) viral gDNA (when present) within individual cells from a variety of sources. Briefly, individual cells within viable cell suspensions undergo encapsulation in the Mission Bio Tapestri platform to form isolated droplets (Figure 1A, boxes 1-2) that contain protease enzymes to allow access to, and thereby release of, viral and host gDNA. The lysate, including the now histone-free DNA, then undergoes a second encapsulation step, wherein the contents are merged with barcoding beads and reagents for target PCR amplification (Figure 1A, boxes 3-5). This approach is described in more detail by Eastburn *et al.*^23^. Amplified regions, or amplicons, for this project consisted of approximately 250-300 base pairs (bp), collectively spanning nearly the entire S[imian]IV genome (Figure 1B), the 2-LTR circularizing junction (Figure 1C), and the following host genes: the SPRY domain of the *TRIM*5*α* macaque gene and *RPP*30 gene (primer details provided in Table S2). Full-genome coverage of SIV could not be attained owing to the repeat of the 5’ LTR (also present as the 3’ LTR) and because the forward and reverse primers for the first amplicon were designed so as to assess 2-LTR formation status (Figure 1B, C). It is important to note that we did not distinguish between remaining DNA forms, which include linear, unintegrated DNA and 1-LTR circle DNA. Host gene primers served to support the cell-calling algorithm of the bioinformatic pipeline, to act as a control for confirming confidence in single-cell capture, and for measuring bioinformatic pipeline performance. See the methods section for more details on the scDNA-seq workflow and strategies for enhancing sample integrity.

**Figure 1.**
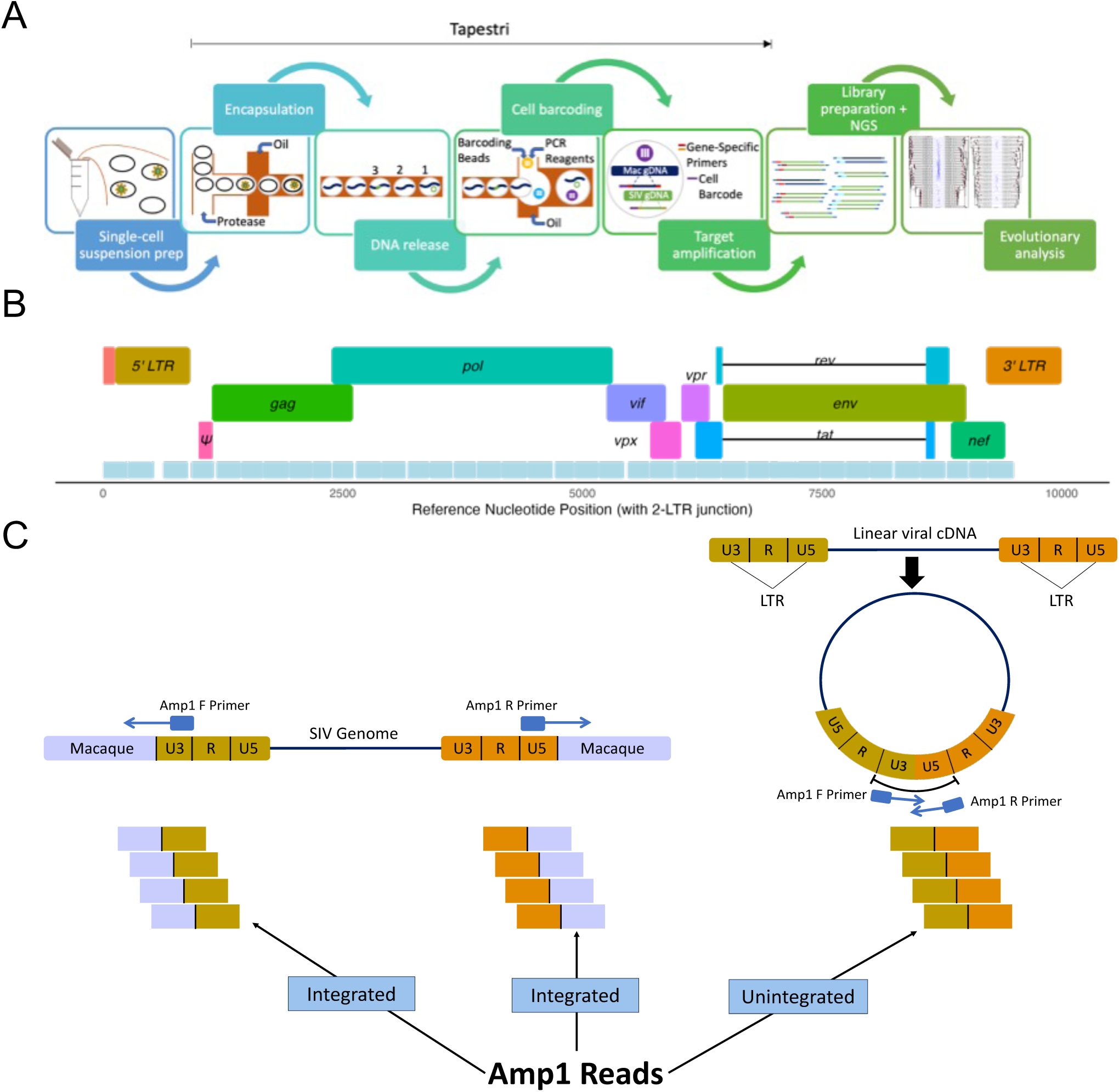
scDNA-seq increases viral genomic sequencing depth and breadth via a droplet microfluidic platform and enables determination of genome integration status. **A)** Schematic overview of the scDNA-seq experimental workflow using the Tapestri® platform adapted for viruses. The encapsulated cell lysates numbered in the DNA release step exemplify the different ways in which scDNA-seq has been designed to capture macaque and viral (SIV) genomic DNA from individual infected or uninfected cells. Following library preparation and next-generation sequencing of resulting DNA, scDNA-seq allows for enhanced phylogenetic analysis and in-depth study of the defective and intact viral genomic landscape on a per sample basis. **B** Amplicon coverage across the SIV reference genome (GenBank accession Available on publication). *Amplicon 1 (Amp1), covering the circularizing junction connecting the 5’ and 3’ untranslated (U5U3) fragments within the long-terminal repeat (LTR) regions of 2-LTR SIV DNA forms. **C)** Visual representation of Amp1 construction and use in distinguishing 2-LTR circularized SIV DNA from linear DNA. Retrieval of complete Amp1 reads indicate virus present in 2-LTR form, whereas Amp1 reads that do not map to the U5U3 junction in the reference sequence are discarded and assumed to be indicative of linear DNA.

Below we describe controlled experiments designed to determine optimal coverage of the designed primers, allowing for reliable characterization of SIV DNA genomic defects within the blood and lymph node compartments and, ultimately, the impact of defective genomes on phylogenetic inference.

## Results

### Animal infection, treatment, and treatment interruption

Full suppression of viral load (VL) below the reliable limit of detection for the in-house quantitative PCR protocol (50 copies/uL) was achieved for all three macaques (HG02, JA41, and JP70) at 50 dpi, and all animals achieved viral rebound to detectable levels by day 195 (Figure S2).

### Depth in viral DNA genome sequencing

The total number of cells retrieved varied across sample, ranging from 1, 047 − 48, 930, which was largely dependent on sample quality and preparation method (discussed in Methods). The number of viral gDNA copies recovered per sample ranged from 1 − 720. Of note, biological replicates (2) were performed for cART LN and post-cART PBMC samples from animal HG02 for comparison of sequencing platforms, resulting in increased number of sequences relative to other samples. The number of total SIV genome copies for PBMCs during cART was low in comparison to post-cART and to that of LN, both during and following cART interruption (Figure 2A). Earlier studies of viral DNA populations in LN and PBMC presented models supporting high similarity between the two populations during cART^37^. However, more recent studies have claimed to detect viral evolution in tissues collected during cART that imply greater differential replication inhibition between anatomical sites^13^. Paired with scrutiny over whether cART can effectively penetrate LN follicles to prevent ongoing viral replication^38^, the increased number of SIV+ LN cells identified during cART by comparison with PBMCs may be a result of exploited putative sanctuary sites within LNs. Of the total 11, 411 SIV genomes recovered, mean coverage was 25.51% of the available genome (2, 423 nucleotides), suggesting a significant presence of replication-incompetent genomes (Figure 2B). On average, the mean coverage was similar between cART (26.4%) and post-cART (24.8%), as well as between LN (29.2%) and PBMC samples (21.5%).

**Figure 2.**
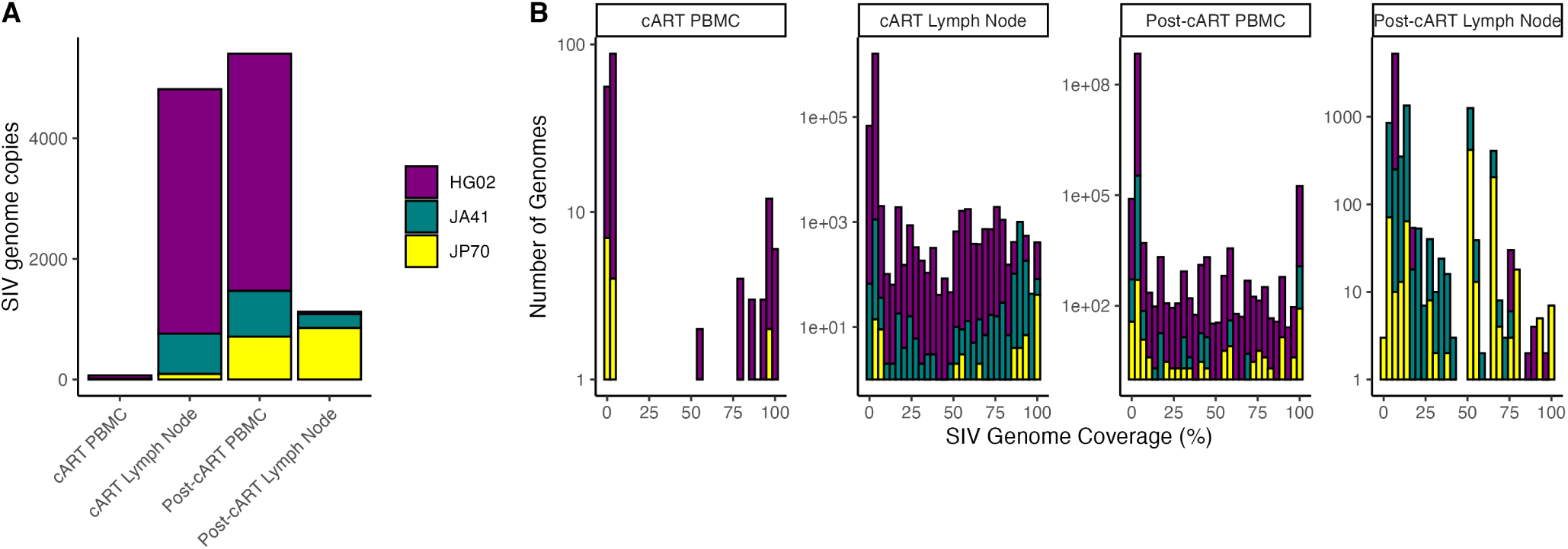
scDNA-seq depth and coverage across collected tissues (lymph node and PBMC) and time points (cART and post-cART) for each macaque. **(A)** Total number of SIV genomes recovered per tissue, time point and animal. **(B)** Distribution of SIV genome coverage per tissue, time point, and animal. Coverage was based on the 9,252-base pair reference genome (2-LTR junction excluded). Colors represent individual animals.

Intra-host genetic diversity estimates for the *env gp120* region were consistent with previous studies, with mean pairwise genetic distances across animals ranging from 1.79*E*-03 to 5.47*E*-03 nucleotide substitutions/site (subs/site, Figure S3)^29,39–41^, though with a select minority of sequences (particularly in post-cART samples) approaching 5.0*E*-02 subs/site, as might be expected with increasing depth of sequencing^42^. However, hypermutation present at sub-detectable levels when using Hypermut 2.0^43^ may also be present (see Methods). The percentage of *env gp120* sequence pairs demonstrating 100% sequence identity was greater within the LN than PBMCs, comprising up to as many as 100% of sequences during cART and following cART interruption for animals JA41 (n=78) and HG02 (n=28), respectively, suggesting a greater presence of clonal amplification within this tissue than in the blood. Up to 55.6% of identical sequence pairs (animal JA41) involved both PBMCs and lymph node, supporting clonal expansion of infected cells in or out of tissue compartments. However, accurate detection of clonal sequences can not be obtained here, as it requires sequencing of integration sites^31,44^.

Sequencing and bioinformatic errors were assessed using *TRIM*5*α* sequence variants (amplicon 42 in Table S2), with the expectation of rare multi-allelic sites at frequencies consistent with heterozygosity. A total of ten nucleotide sites were shared across all animals at which more than one allele was observed (Figure S4). Only one instance of a third allele was observed and at a frequency of 0.006%. Two of the ten sites (positions 29190 and 29186) presented with minor allele frequencies of < 1%, potentially a consequence of sequencing errors specific to the Illumina platform (e.g., certain trimer motifs, such as GGT and AGT, have been associated with Illumina machines^45^). Both trimers at these positions were represented by majority AGG; however, only one presented with a known problematic motif (AGT) in the reference sequence. The other minor allele trimer (reference motif AGC), while not established as a top ten error trimer for Illumina, remained suspect as a platform error, as it occurred in all three animals with a maximum incidence of 0.1%. Approximately 69% (18/26) of multi-allelic sites harbored minor alleles, though present at < 1% (Figure S5), indicating a relatively low error rate of 0.17% (proportion of cells containing at least one dominant sequencing error) generated during sequencing and/or preparation of sequence library. Results thus demonstrate the reliability in calling variants for consensus sequence reconstruction for the vast majority of cells present. Taken together, these results validate earlier efforts demonstrating that this scDNA-seq approach enables the retrieval of a substantial number of sequences while keeping a low estimated error rate.

### Inference of viral DNA replication competency

We next explored whether this scDNA-seq approach can reliably classify viral genomes based on defects. Digital Droplet PCR (ddPCR), designed to detect SIV regions non-overlapping with scDNA-seq amplicons (Figure S6), was used to corroborate scDNA-seq coverage of the SIV genome. ddPCR primers and probes were adapted from HIV- and SIV-specific primers and probes, as well as an additional probe targeting the RPP30 gene, used in Bender *et al.*^2929^. Adaptation was undertaken so as to increase coverage across the SIV genome relative to previous applications, as well as to increase permissivity to hypermutant sequences. SIV-specific primers and probes were tested on WT plasmid and a hypermutant plasmid generated using site-directed mutagenesis (SDM) of all possible GG|GA dinucleotides in the WT strain, representing APOBEC-mediated hypermutation. See methods section for more details on plasmid construction and transfection. Equivalent relative abundance of probes covering the 5’ and 3’ regions of the polymerase (*pol*) gene, packaging signal (*psi*) region, and 3’ region of *env* (> 99.3%) was observed compared with 5’ *env* probe, indicating ideal performance for both WT and hypermutant plasmid sequences. Following 24-hour transfection of human embryonic kidney HEK293T cells with each stock plasmid, cell suspensions were frozen for additional ddPCR analysis. ddPCR revealed a >1:1 plasmid:cell ratio (mean of 2.64 and 2.05 plasmids per cell), represented by the RPP30 probe, for WT- and hypermutant-transfected cells, respectively. The same cell aliquots were then used in scDNA-seq.

A 100% transfection rate of HEK239T cells (total of 3,772) with WT plasmid was similarly determined using scDNA-seq, though with non-uniform coverage ( Figure S7). Consecutive amplicon dropouts (up to three) were observed. To account for this dropout, loss of four consecutive amplicons was considered the criterion for classification of genomes as harboring at least one large internal deletion. With coverages of 99.5% and 98.3% in the 5’ and 3’ flanking amplicons, respectively, loss of a single flanking amplicon was considered the final criterion for classification of a genome as 5’ and/or 3’ deletion-defective. Based on these criteria, 97.16% of WT-transfected cells were classified as containing intact sequences (Table 1). Remaining false positive defective virus were observed as deletions in (*psi*) or stop codon mutations (0.21%), large deletions in the 5’ and/or 3’ ends of the genome (2.45%), or as hypermutations (0.19%). These defects may be the result of mutations during PCR amplification in the viral genome or in the cell barcode used in cell calling in the bioinformatic pipeline. The 2-LTR junction (Amp1) was detected in 0.16% of cells, which may represent virus still in plasmid form.

**Table 1.** Pipeline classification of HEK293T cells transfected with wild-type SIV plasmid. Results are reported in percent of cells (*n* = 3772) identified. *Adjusted for amplicon dropouts.

| Classification | % of cells |
| --- | --- |
| Intact | 97.16% |
| Psi deletion or point mutation | 0.21% |
| 5' Deletion | 0.19% |
| 3' Deletion | 1.54% |
| Large internal deletions (after adjustment*) | 0% |
| 3' + 5' Deletion | 0.72% |
| Hypermutated | 0.19% |

PCR-mediated SDM was used to generate hypermutant plasmids; however, 100% accuracy with SDM can be challenging, especially with AT-rich genome templates such as retroviruses^46^. Hence, it was not surprising to find that hypermutant plasmid sequences resulting from scDNA-seq comprised a wide range of hypermutation (1-40% possible sites, Figure 3A). As a note, these values likely represent an underestimate because they were only calculated for sequencing reads present and do not account for additional hypermutations resulting in amplicon dropout. All hypermutant sequences, regardless of their level of hypermutation, were classified as either hypermutant (according to Hypermut v2.0^43^) or putatively intact, with small or no internal deletions, as described above (Figure 3A). The absence of deletion defects within the control plasmid may not translate to biological hypermutation variation, however. Hypermutated sequences may falsely contribute to deletion-defective category numbers if remaining sequenced regions are not detected as significantly hypermutated by Hypermut. Contrary to WT-transfected cells, the 2-LTR junction was not detected among hypermutant plasmids (Figure 3B). The remainder of amplicons were present in at least one sequence, with certain amplicons demonstrating an ability to withstand more hypermutations in primer-binding regions than others (Figure 3B). Amplicons 8 (*gag*) and 30 (*env*) were present in 100% and 99.2% of the sequences, respectively, and regardless of level of hypermutation. Each of these regions were permissive to the maximum of two hypermutated dinucleotide sets in the forward primer sites. Hence, >99.2% of intact hypermutated sequences present in a given sample should be detected using this scDNA-seq method using the existing criteria of the presence of at least two amplicons (100% if relaxed to a single amplicon). The highest reported level of hypermutation was observed in amplicon 3, which permitted up to 3 hypermutated dinucleotides, though 4 sets are possible here, potentially explaining its loss in 59% of sequences.

**Figure 3.**
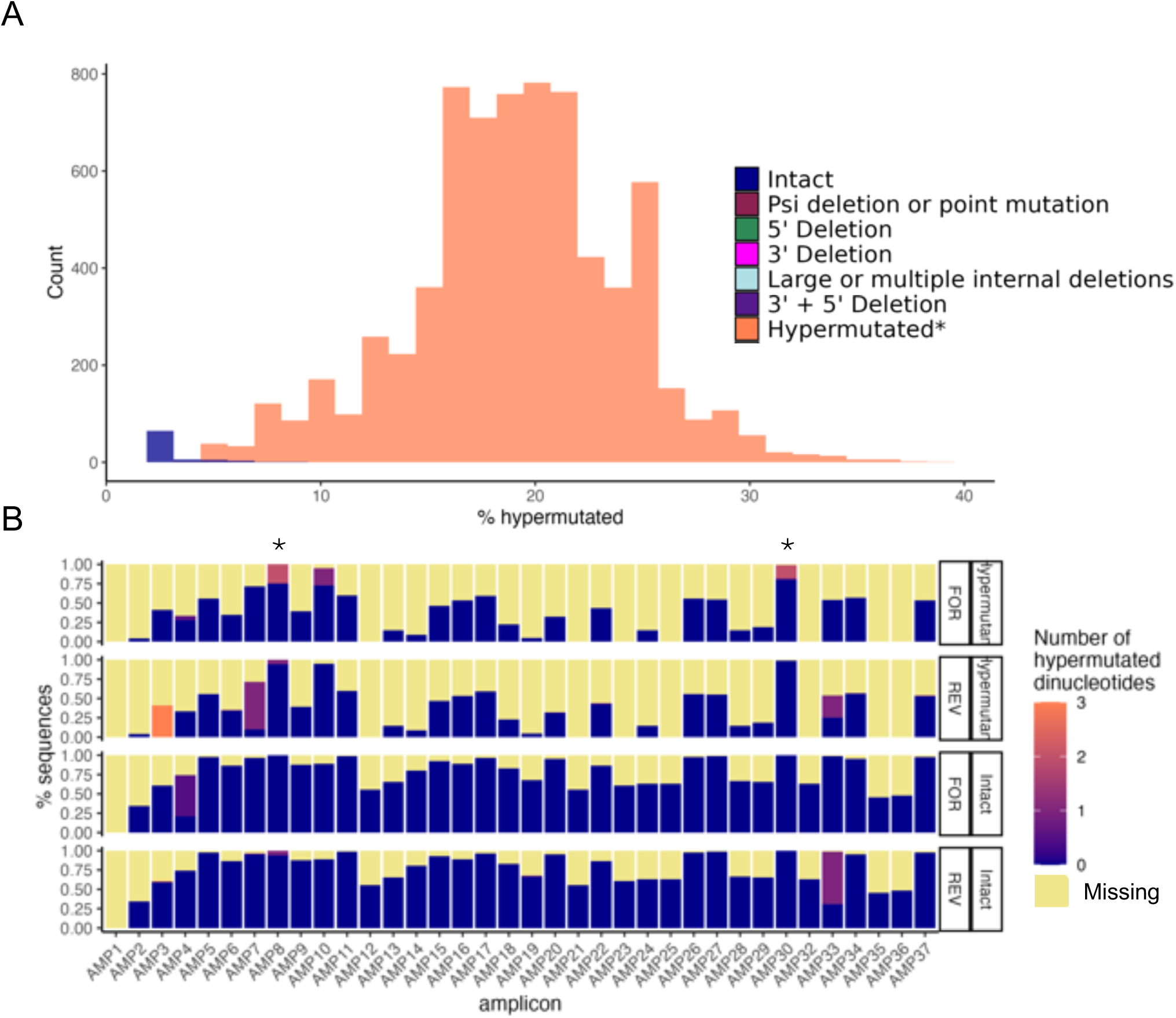
scDNA-seq application to HEK293T cells transfected with hypermutated SIV plasmid. **(A)** Distribution of sequences according to % of hypermutated sites within sequenced regions, demonstrating range of hypermutation resulting from SDM. Sequences were grouped according to classification using the scDNA-seq pipeline (legend at right). **(B)** Sequence classification representation using the scDNA-seq pipeline (legend at left). **(C)** Range of number of hypermutated dinucleotides (GG|GA) present within primer-binding locations for sequences classified as hypermutated (top half) or intact (bottom half) using the scDNA-seq pipeline. *Amplicons present in >99% of sequences. Amplicons missing entirely from final sequences are represented in yellow. FOR=forward primer, REV=reverse primer.

Following plasmid validation, PBMC samples taken at necropsy from the three animals were similarly used in a ddPCR and scDNA-seq comparison. Specifically, ddPCR-probed regions were evaluated among resulting scDNA-seq sequences for completeness (>90% non-ambiguous sites considered complete), and the fold difference from ddPCR was measured (in percent positive cells). A < 0.25-fold difference was observed, with the exception of probes located in the 5’ and 3’ *env* gene regions (Figures 4A,B). For these regions, scDNA-seq resulted in nearly 4-fold greater copy numbers than ddPCR in animal HG02 (Figures 4A,B). Upon examination of mutations, insertions, and deletions in primer- and probe-binding regions within the sequences, reduced detection with ddPCR could be explained by >4 nucleotide deletions in the 5’ *env* reverse primer and 3’ *env* forward primers, not present in any other region or animal (Figure 4C), demonstrating the effectiveness of the scDNA-seq approach in capturing these given regions over ddPCR for these particular samples.

**Figure 4.**
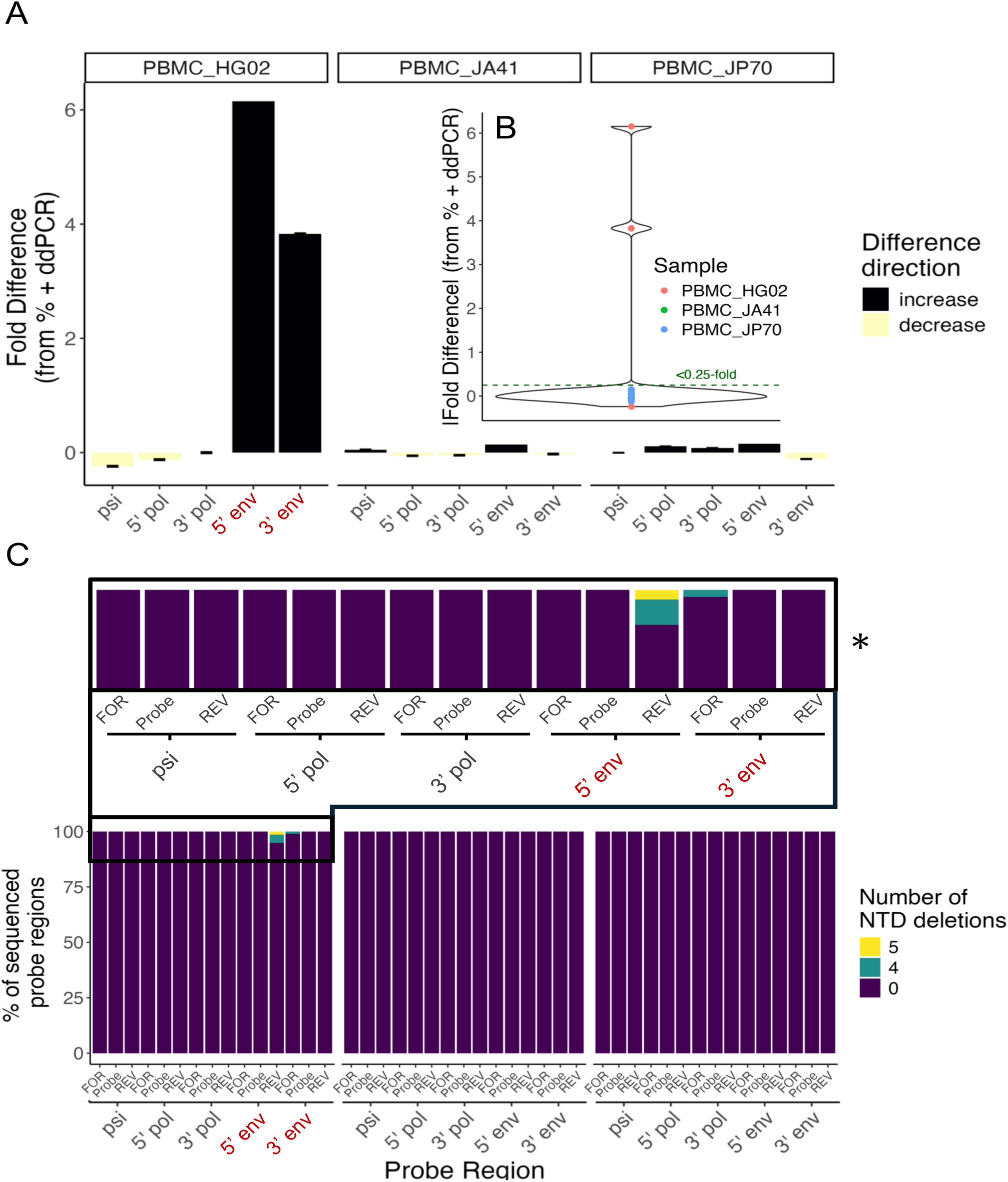
Comparison of scDNA-seq with ddPCR for PBMC samples. **(A)** Difference in detection of five probed regions between ddPCR and scDNA-seq. ddPCR-probed regions were evaluated among resulting scDNA-seq sequences for completeness (>90% non-ambiguous sites considered complete), and fold difference of scDNA-seq from ddPCR was measured in percent-positive cells. **(B)** Fold difference in scDNA-seq from ddPCR, expressed in absolute value. **(C)** Number of nucleotide (NTD) deletions present among forward (FOR) primer, reverse (REV) primer, and probe sites for each of the five ddPCR-probed regions. *Inset emphasizing the only animal and regions comprising nucleotide deletions, corresponding with large fold-differences observed in A and B.

When genome classification was applied to animal sequences, animal-specific patterns were evident, with intact sequences comprising a wide range of 0.4 − 20% of the viral DNA population (Figure 5). Assuming all putative intact virus are replication-competent, the greater relative abundance of intact virus in animal JP70 may explain the more rapid rebound to greater viral load levels in this animal following therapy interruption (Figure S2). With regards to defects, an abundance of putative deletions in either or both of the 5’ and 3’ genomic ends in all animals over internal deletions, point mutations, and hypermutation, was evident. The largest pairwise genetic distances were observed among these deletion-defective variants, rather than intact virus, suggesting remaining genome regions may harbor mutations considered informative for phylogenetic inference.

**Figure 5.**
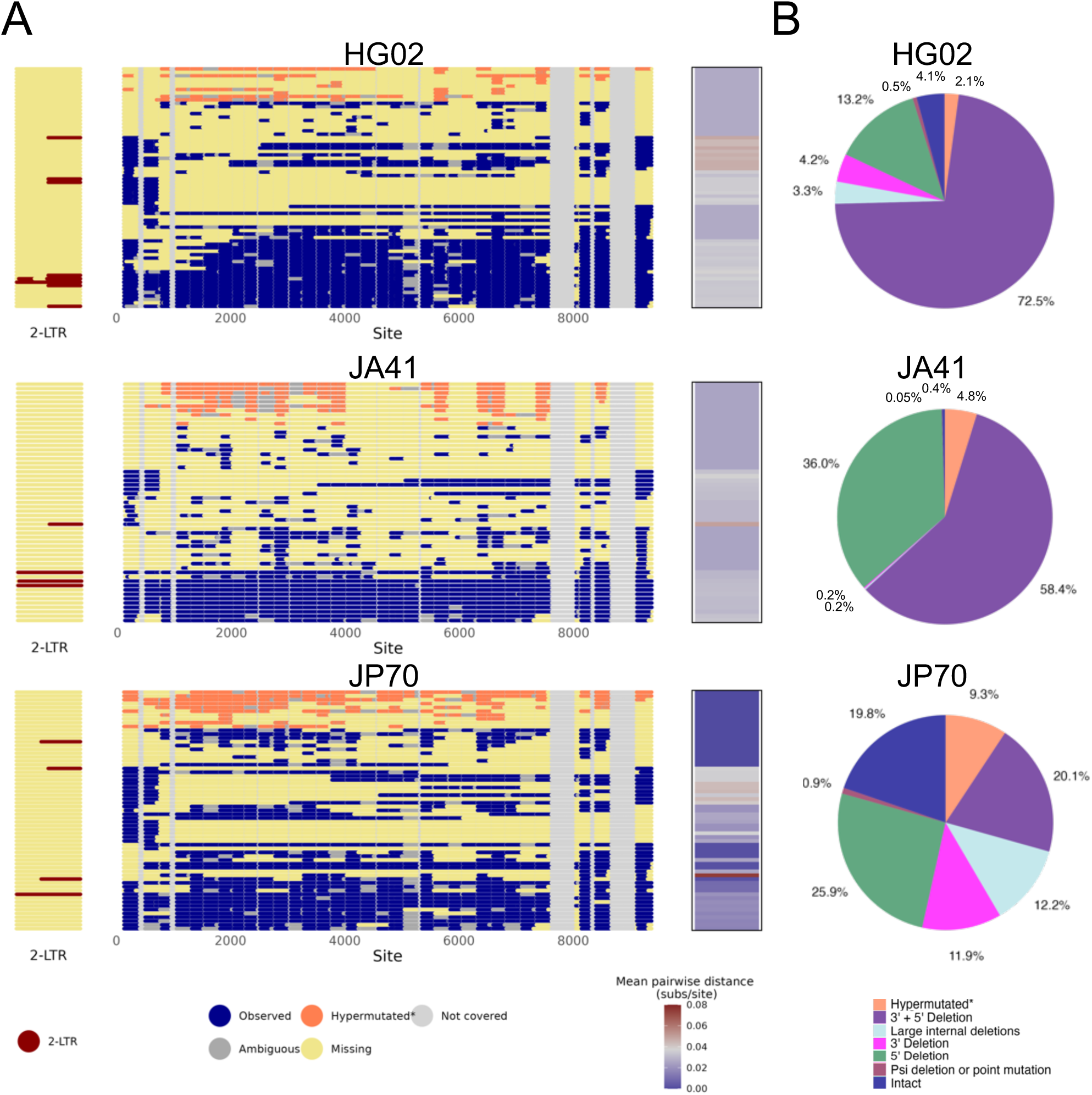
Defective profiles of SIV DNA genome sequences detected using scDNA-seq for each animal. **(A)** Representative sequences from defect classification categories (Table 1. For each category, up to 10 sequences were chosen as representatives based on the sum of pairwise genetic distances when compared to remaining sequences in the corresponding category. Bases are colored according to presence within the 2-LTR junction (maroon) and remaining genome (blue), or absence in the form of ambiguity (grey), gap (yellow), or lack of coverage with existing primers (light grey). *All non-ambiguous bases for sequences determined to be hypermutated have been colored (orange) for easier identification of hypermutated sequences. Mean pairwise distances, scaled in substitutions/site, are also graphed (right), providing a degree of representation. **(B)** True defective class representation among all sequences for each animal (legend at bottom).

### Phylogenetic resolution

To investigate improvements in the reliability of viral evolution reconstruction using genomes generated from the scDNA-seq method, we first compared the phylogenetic signal of NFL genomes with their respective *gp120* fragments (i.e., NFL genomes obtained by scDNA-seq were disaggregated into subgenomic *gp120*) (Figure 6 and Figure S9). Only those genomes with intact (*≥* 90%) *gp120* were thus included for this analysis, representing a comparison of scDNA-seq with traditional *gp120*-targeted single-genome amplification and Sanger sequencing^47^. Sequences identified as hypermutant underwent masking of potentially hypermutated sites, as in^48^ (see Methods for additional details). Likelihood mapping was used to visualize phylogenetic signal, which provides a high-level summary of the percentage of randomly sampled sequence quartets that could be reliably resolved in the form of one of three possible bifurcating phylogenies (represented by the corners of the triangles in Figure 6A) compared to the percentage unresolved (center of the triangle). Animals exhibited up to a nearly 5-fold reduction in the number of unresolved phylogenies for NFL genomes compared with their *gp120* counterparts (Figure S8).

**Figure 6.**
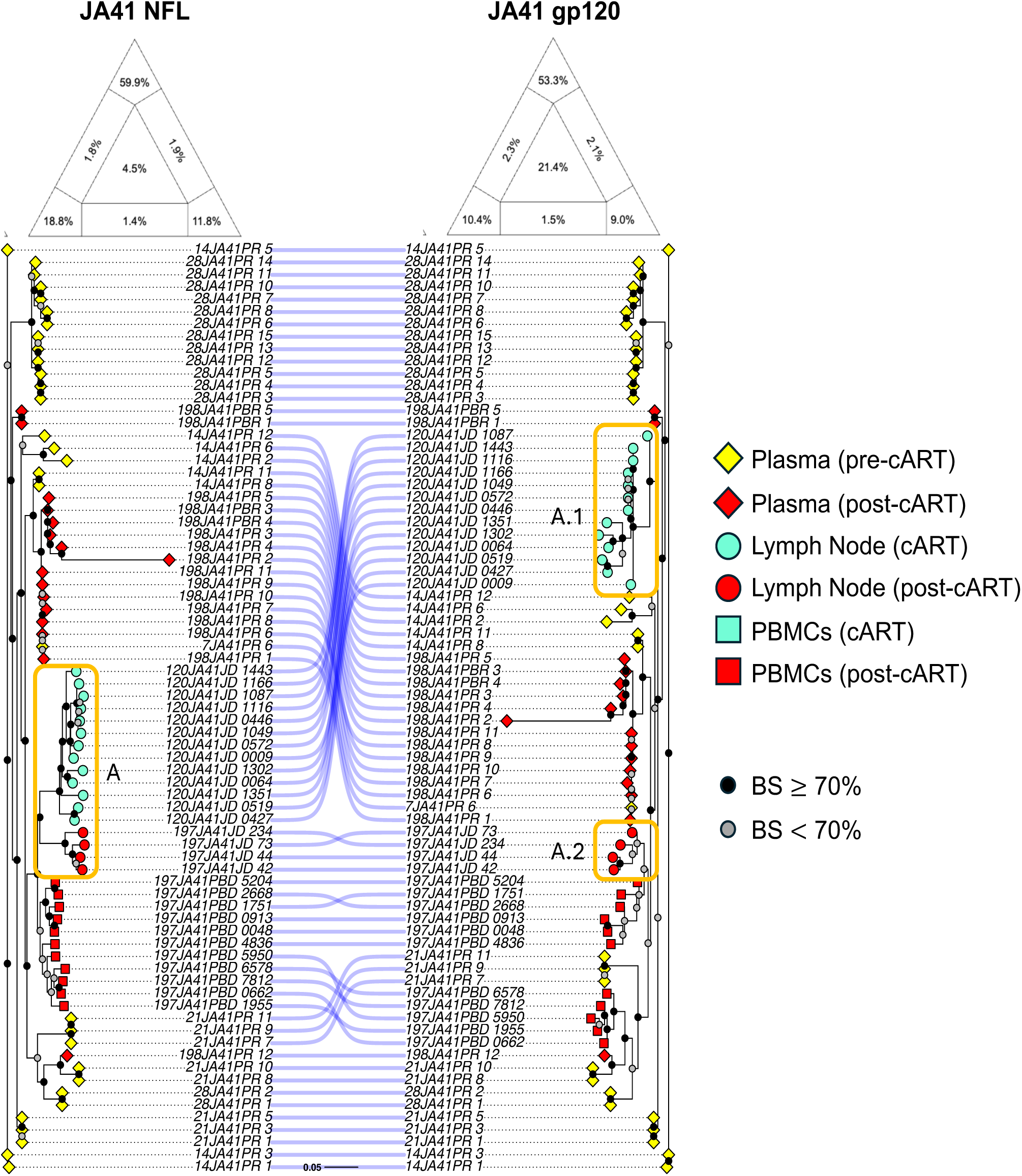
Comparison of phylogenetic resolution and placement for full genomic sequences (left) and corresponding fragments restricted to subgenomic *env gp120* (right) for animal JA41. Phylogenetic likelihood maps (top) demonstrate the percentage of sampled sequence quartets classified as phylogenetically resolved (corners), partially resolved (outer edges), or unresolved (center) during phylogenetic reconstruction. Maximum likelihood phylogenetic trees for HG02 samples were analyzed using scDNA-seq (legend at right). Shapes within the tree indicate tissue type (square: PBMC, circle: LN, diamond: plasma), whereas colors indicate time point (yellow = pre-cART, cyan: cART, red: post-cART). Support for branching events was provided through the transfer bootstrap expectation^49^, with *≥* 70% considered well-supported and represented by a black circle and < 70% represented by a gray circle. Asterisks indicate genomes classified as intact. Clades A (full genomic sequences on left) and A.1/A.2 (*gp120* sequences on right) represent one example of sequences with different branching patterns between the two trees.

Maximum likelihood trees were then reconstructed for NFL and *gp120* datasets above, including also pre-cART and post-cART plasma viral *gp120* RNA sequences, for a quantitative description of tree topological differences (Figure 6B and Figure S9). Robinson-Foulds (RF) distances between the two trees for each animal were first calculated in order to describe quantitatively the differences in topology (branching patterns)^50^. Briefly, for two trees with identical sets of leaves, the RF distance describes the number of non-trivial bifurcations that differ between trees. Final values were normalized by the maximum potential RF distance. RF distances ranged from 0.44 (JA41) to 0.71 (HG02) (Table S3), indicating the NFL and *gp120* trees were, at best, 56% away from completely randomized (100%) tree topologies.

Differing topologies translate to the potential for differences in ancestral sequences and/or states (e.g., tissue) within the tree. As described in the introduction, ancestral state reconstruction within a phylogeny can be used to infer state transitions and thus origins but assumes that the individual phylogenetic relationships within the tree are known, or at least reliable; this is not always the case, as can be seen from the percentage of sequences considered unresolved in Figure 6 and differences in tree topology (Table S3). Increased resolution has the potential to result in differing branching patterns altogether as compared with lower-resolution data. In the JA41 NFL genome tree, for example, a subtree of closely related post-cART LN sequences exhibited shared recent common ancestry with LN sequences sampled during cART, which was supported in *≥* 70% of bootstrap replicates^49^ (Figure 6B, clade “A”). Contrastingly, shared common ancestry was split in the *gp120* tree (Figure 6B, “A.1” and “A.2”), suggesting differing origins could be inferred in the absence of the additional sequence information provided from the NFL genomes.

Following the comparison relying on complete *env*, we incorporated additional inclusion scenarios in likelihood mapping analysis, comprising 1) all NFL genomes generated from the scDNA-seq approach and genomic regions contained within and 2) only those genomes considered to be likely intact and lacking significant hypermutation, stop codon-altering mutations, and large internal deletions. For the first scenario, hypermutant sequences were included following masking of hypermutated sites, as described above. Intact genomes appeared to generate a further reduction in unresolved quartets relative to NFL genomes containing intact *gp120* (Figure S8). Contrastingly, the largest percentage of quartets considered to be unresolved belonged to the datasets including all NFL genomes, ranging between 45 − 92.5%). This finding was not surprising, given that overlap of genomic regions between sampled quartets is under-represented as with intact genomes or genomes containing intact *gp120*. A more reliable estimate of contribution of this unique dataset type to phylogenetic resolution would perhaps be attainable with a likelihood mapping approach that is restricted to only those quartets with genomic overlap. Without this filtering, we can only conclude that phylogenetic resolution is enhanced from traditional *gp120*-focused sequencing with the addition of genomic regions outside *gp120*, regardless of integrity, though tree reconstruction and downstream inferences may be burdened by noise with the inclusion of a large number of genomes that do not overlap in sequence.

### Phyloanatomic inference

We next sought to more rigorously evaluate differences in inferred tissue origins in order to determine if informative phylogenetic relationships were still being formed despite the addition of phylogenetic noise. Transitions between tissue states along well-supported branches were identified for the differing tree scenarios above: 1) *gp120* only and 2) NFL genomes containing >90% *gp120*, as in Figure 6B, but also for 3) intact-only sequences and 4) the full suite of NFL sequences provided from the scDNA-seq approach, including masked hypermutants. An illustration of representative sequences in each of these four categories can be found in Figure 7A. Pre-cART and post-cART plasma *gp120* RNA sequences were again included here. Support for branches was considered in the form of the transfer bootstrap expectation (TBE)^49^, and only well-supported branches were used in the identification of tissue state transitions. Similarly, only inferred states with scaled likelihood > 0.90 were considered. Importantly, all tree types indicated plasma pre-cART origins for every cART and post-cART tissue, consistent with rapid dissemination of virus into tissues during acute infection and establishment of a reservoir of infection. However, overall, trees containing either intact-only or *gp120*-only sequences produced the fewest supported transitions between tissues and the highest transition rate differences between tissue pairs (Figure 7B). The greatest number of supported transitions among tissues and time points was consistently reported for trees containing all NFL genomes resulting from scDNA-seq, regardless of their integrity. Transition rates among tissues and time points within these trees were also more evenly distributed, placing less emphasis on seeding of tissues during acute infection (pre-cART). Furthermore, unlike the trees reconstructed using the stricter (and not necessarily justified) inclusion criteria, there exists increased support for mixing of the viral population between tissues during cART when all genomes are considered, indicating a significant contribution of incomplete genomes to statistical phyloanatomic inference. In other words, while a large proportion of these incomplete genomes are not readily resolved among randomly sampled sequence quartets, inclusion of these genomes strengthened support within areas of the tree involving shared ancestry between differing tissues.

**Figure 7.**
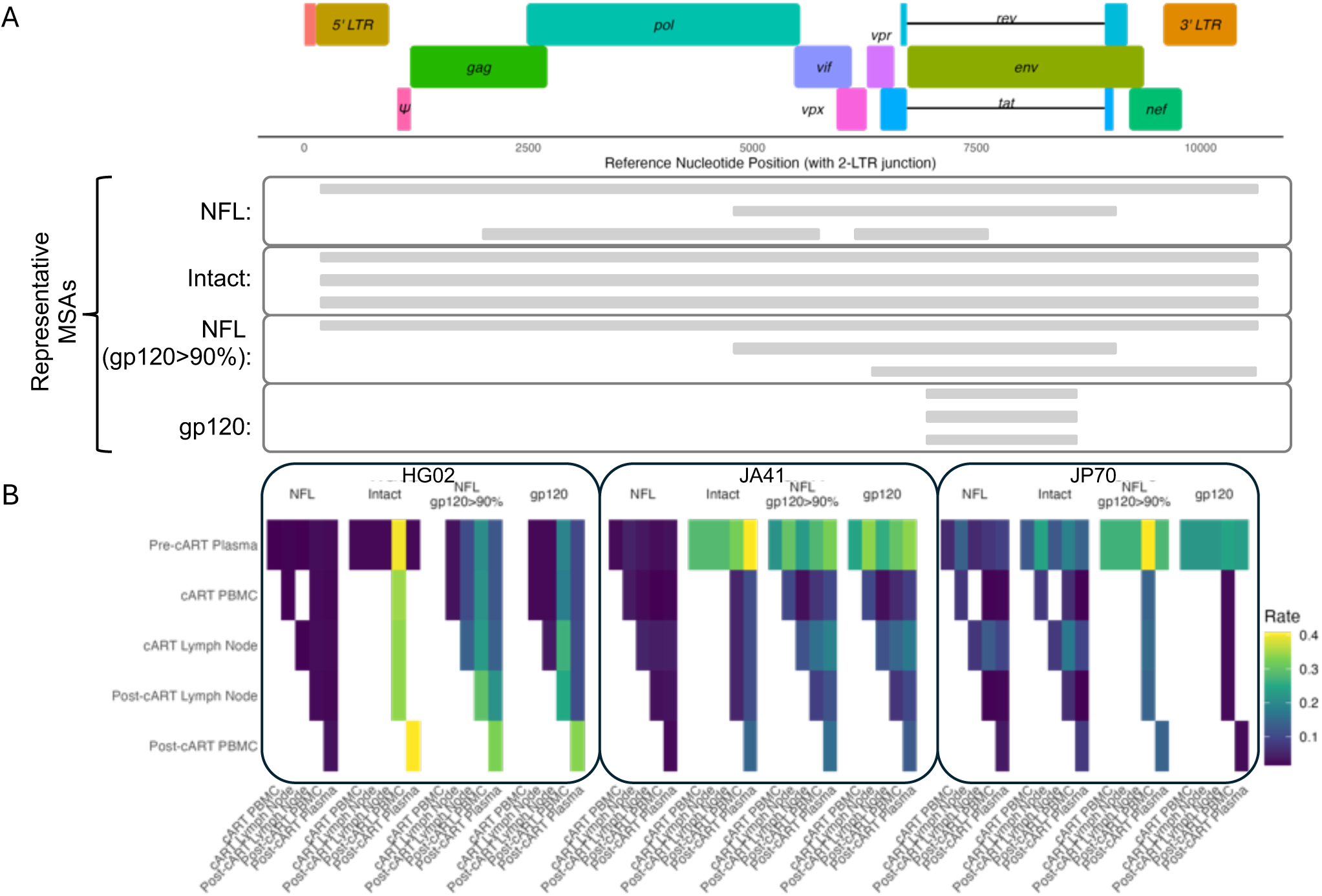
Phyloanatomic comparison among trees inferred from differing multiple sequence alignment (MSA) inclusion criteria. **A)** Representative MSAs for each of four inclusion scenarios. Near-full-length (NFL) genome sequences correspond to all DNA sequences resulting from the scDNA-seq workflow. Intact sequences are determined based on absence of large internal and flanking deletions, as well additional deleterious mutations. NFL genomes with *≥* 90% *gp120* coverage are considered in their entirety in the third category. Finally, NFL genomes with *≥* 90% *gp120* were stripped to the *gp120* region only as the fourth category. **B)** Transitions between tissue and time point class (e.g., PBMCs during cART) for each animal and scenario. Transitions were determined using the inferred ancestral states along branches within the animal-specific phylogeny (e.g., Figure 6). Only well-supported branching events (> 70% TBE support) and states with scaled likelihood > 0.90 were considered.

## Discussion

Existence of the HIV reservoir, adept to replenishing viremia upon therapy interruption, has continued to pose a major barrier towards attaining HIV eradication within its host. Uncovering critical reservoir dynamics that facilitate the long-term maintenance of HIV are imperative to more precisely target the tissue(s) responsible for viral rebound^51^. Several sequence-based methods have been developed to further investigations of viral DNA populations, though often limited by the trade-off in depth and breadth, as well as the ability to correctly classify integration of the viral genome^26,52–55^. Previous assays that have successfully incorporated targeted, near-full-length (NFL) genome sequencing of HIV/SIV have been devised with the replicating virus in mind, incorporating a limited number of amplicons that ultimately restrict the size and coverage of the defective population pool^7,10,30,31,56^. This restriction represents a missed opportunity for inclusion of mutational histories that improve phylogenetic inference. Here, we describe the use of single-cell DNA sequencing (scDNA-seq) to target a large number (37) of relatively conserved regions in the SIV genome in order to determine the level of enhanced resolution provided by the defective genome population for improved reconstruction of viral reservoir dynamics across tissues in a treatment interruption macaque model.

Through controlled sequencing and complementary droplet digital quantitation, we demonstrated herein the capability of the scDNA-seq approach to reliably distinguish intact and defective viral genomes. DNA sequences collected from PBMCs and lymph node were characterized according to defect for three animals during cART and subsequent cART interruption.

Among all three animals, deletions encompassing either one or both ends of the SIV genome comprised the dominant defect classification, corroborating existing evidence that deletions are the most common defect within the HIV DNA population^10,57^. Importantly, however, these defective genomes harbored extensive genetic diversity that prompted investigation into their ability to aid in phylogenetic reconstruction and downstream inferences, such as tissue migration.

Though many of the genomes produced from the scDNA-seq method can contribute noise to the phylogenetic reconstruction process owing to large deletions, we show here that inclusion of incomplete DNA genomes is critical for recovery of viral migration patterns. For example, in two of the three animals, the use of intact- or *gp120*-only sequences limited the origins of plasma viral rebound to post-cART PBMCs or pre-cART plasma, providing no information regarding anatomical reservoir locations during therapy. The increased representation of well-supported transitions in space and time, relative to the phylogenies reconstructed from *gp120* or intact virus only, demonstrated not only tissue origins of rebound virus but offered evidence of movement of virus or virus-infected cells between tissues during therapy. It is important to keep in mind, however, that this inclusion of noise increases the need for reliance on well-supported branches. Movement across anatomical compartments poses issues for quantitative analysis of reservoir dynamics, such as estimation of the tissue-specific half-life for feasibility of viral eradication. More importantly, quantifying and distinguishing movement resulting from cellular proliferation and trafficking versus cell-free migration is imperative for the design of more targeted interventions. Further phyloanatomic investigation using statistical approaches applied to more available tissues from the macaque model are needed to not only determine the full extent to which inclusion of the defective genome population improves migration inferences but also to disentangle the distinct contributions of cellular and viral activity to the spread of virus among tissues during therapy.

We recognize that, regardless of our increased resolution in defective classification and characterization owing to shorter amplicons, targeted sequencing has its own limitations. As is often observed for heterogeneous viruses, mutations in primer-binding sites can result in sub-optimal primer binding “amplicon dropout” (observed in Figure S7). This heterogeneity can exist in the form of divergence from the reference sequence or intra-host population heterogeneity. We recognize that, using this approach, we were unable to fully distinguish regional deletions from amplicon dropouts. Isolated, internal amplicon dropouts were not observed among the macaque samples; however, even multiple amplicon dropouts within the flanking regions of the genome deserve additional attention and may have resulted in the underestimation of the number of intact genomes. Improvement of potential vulnerabilities in the original primer design may be found with advancements in long-read sequencing or targeted wobble-base replacement.

In the face of the above-described limitations, the scDNA-seq approach permitted unparalleled depth of NFL viral DNA genome sequencing that serves to (1) broaden analyses of not only intact, but defective viral populations that persist during cART and (2) improve confidence in phyloanatomic patterns. Now armored with multi-omic capabilities^31,58^, this approach offers potentially expansive and informative insight into the maintenance and stability of the reservoir populations and underlying cellular compositions.

## Methods

### Ethics statement

All animal procedures were performed by the Tulane National Primate Research Center (TNPRC) in accordance with Tulane University’s Institutional Animal Care and Use Committee (IACUC) protocol P0376. The macaques were housed in groups or pairs at Tulane animal facilities accredited by the Association for Assessment and Accreditation of Laboratory Animal Care (AAALAC). Details surrounding animal welfare including environmental parameters and standard practices for treatment of non-human primates in research were followed as outlined in the *Guide for the Care and Use of Laboratory Animals*^59^. All possible measures were employed to minimize discomfort of the animals. Once IACUC defined endpoints were reached, macaques were humanely euthanized following the standard method of euthanasia for non-human primates at the TNPRC, consistent with the recommendations of the American Veterinary Medical Association Guidelines on Euthanasia.

### Study population

Two Indian *Rhesus* macaque (*Macaca mulatta*) groups, referred to in the present study as groups 2.1 (n=4) and 2.2 (n=4), were originally staggered for use, representing a single cohort of SIV-infected, cART-interrupted animals. The study cohort was intravenously inoculated with the same SIVmac251 viral swarm at a dosage of 100 TCID50^60,61^. Following 35 days post-infection (dpi), at which median viral set point was established, cART treatment was initiated. The cART regimen comprised of two reverse transcriptase inhibitors: Tenofovir, 20 mg/kg (Gilead) and Emtricitabine, 30 mg/kg (Gilead) administered daily by subcutaneous injection, and one integrase inhibitor: Raltegravir, 22 mg/kg (Merck) twice daily by oral administration. Drug dosages were based on a protocol developed by Dr. Kenneth Williams’ lab at Boston College, Boston, MA, and recommendations by the senior director of Biology at Gilead Sciences, San Francisco, CA. One macaque from group 2.2 was found deceased at 42 dpi and was therefore not included in further longitudinal sampling. At 180 dpi (145 days of drug treatment), cART was suspended to allow viral rebound and disease reemergence. The animal cohort was followed for three additional weeks post-cART and subsequently euthanized for supplemental sample collection. Three animals were chosen from this cohort for evaluation of the scDNA-seq approach based on the increased availability of samples, accounting for the potential need for protocol optimization, as described in more depth below. Of note, references to individual sequences within the text belonging to LN samples contain the designation ’J’ owing to previous sequence naming used by our group^21,60,62^, allowing for easier analytical comparison with previously collected LN samples.

### Viral load quantification

SIV viral load quantification was performed on plasma collected from the three animals at different collection time points prior to, during, and after interruption of cART over the course of 211 days (Figure S2). Viral RNA was extracted from using the QIAamp Viral RNA Mini Kit (Qiagen, Germantown, MD) according to the manufacturer’s instructions. RNA was reverse transcribed using the High-Capacity cDNA Reverse Transcription Kit (Applied Biosystems, Waltham, MA) according to the manufacturer’s instructions. Quantitative PCR was performed using TaqMan^TM^ Fast Advanced Master Mix (ThermoFisher Scientific, Waltham, MA) using 0.4 µM of each of the following primers - 5’-AATTAGATAGATTTGGATTAGCAGAAAGC-3’ (Forward)) and 5’-CACCAGATGACGCAGACAGTATTAT-3’ (Reverse) - and 0.25 µM of the following TaqMan(R) probe - 5’-CAACAGGCTCAGAAAA-FAM-3’ - which target a region near the 5’-end of the group antigen gene (*gag*), based on SIV isolate 239 (M33262.1)^63^. PCR conditions are described in Table S1. The nominal copy numbers for each samples was determined by interpolation onto a standard curve of RNA standards (duplicate reactions for 1:10 dilutions), the limit of detection limit of which was 50 copies/µL.

### Sample collection

A baseline blood draw prior to SIVmac251 inoculation was taken from each animal. Routine weekly physical examinations over the course of pre-infection through necropsy, were accompanied by blood draws for the purpose of monitoring SIV viremic levels and viral reseeding into the peripheral blood. Peripheral lymph node collections were completed on the following days/day ranges post-infection: 14, once between 49 − 63, 120, and 180. Following cART interruption, macaques were euthanized and further sampling obtained at necropsy (197 dpi)) for the previously mentioned sample types. All group 2.1 sample time points preceding 180 dpi were frozen in a homemade freeze media containing 90% FBS : 10% DMSO using Mr. Frosties. At 180 dpi and all time points thereafter, samples were cryopreserved in cryovials to augment storage for downstream live cell recovery. Group 2.2 samples were frozen in the aforementioned live freeze media, and subsequently switched to cryopreservation beginning at 120 dpi.

### Plasma viral RNA isolation and sequencing

Viral envelope *gp120* RNA was isolated at 14, 21 and 28 (pre-cART), and again at 198 dpi (pos-cART interruption). Viral isolates were sequenced using the limiting dilution nested PCR amplification and Sanger sequencing approach described previously in Strickland *et al.*, 2012**^?^**.

### Custom amplicon panel design

A custom reference genome was designed to obtain NFL coverage of the SIVmac251 population, including one amplicon targeting the U5U3 circularizing region (Figure 1C). The SPRY domain of the macaque *TRIM*5*α* gene, as well as the *RPP*30 gene, were added for amplicon coverage inclusive to all cells to aid in examination of pipeline performance for correctly calling variants. Virus amplicon coordinates of the custom reference genome can be located in Table S2.

### Single-cell suspension preparation

Samples containing 1 mL cell suspensions were quickly thawed in a 37°C water bath and carefully aspirated and dispensed to a 1.5 mL microfuge tube. All cell suspension transfers and washes proceeded with the use of wide bore pipette tips to minimize cellular damage. The original storage vial was rinsed with 500 µl of room temperature DPBS (Corning, 21-031-CV) to recover residual cells and gradually transferred to the 1.5 mL microfuge tube. The sample was then centrifuged at 300 *xg* for 5 minutes at 25°C to allow removal of storage media from the cells. The supernatant was discarded and the cell pellet was resuspended in 1 mL of DPBS and subjected to centrifugation as previously described. Following removal of the supernatant, the pellet was carefully resuspended in 50 µl, or as low as 35 µl, of Cell Buffer (Mission Bio) until no clumps were visible then stored on ice. A cell count was performed on the suspension using trypan blue dye exclusion to determine percent viability and quantify live cells by visual inspection. In short, depending on concentration, cell suspensions were diluted with trypan blue either 1:4 or 1:10, then 18 µl of the diluted mixture was pipetted into the loading chamber of a Nexcelom hemacytometer (cat. no. CP2-002). Total number of cells, live and dead, were counted in four corner quadrants using a brightfield microscope with 20X magnification. Counts were averaged and multiplied by dilution factor times 10,000 to obtain cells/mL. The cell suspension was then diluted using Cell Buffer to a concentration of approximately 3,500 cells/µl, and subsequently available for use with the Tapestri® instrument. To circumvent variation in sample integrity as a result of adjustments in storage media and preparation prior to thawing, a modified protocol was implemented to enhance viable cell recovery and executed when necessary (see low-integrity sample enhancement workflow).

### Low-integrity sample enhancement workflow

In circumstances where viability fell below the 80% threshold recommended by Mission Bio for single-cell DNA sequencing, an optimized sample preparation protocol was performed. The workflow utilized the MACS Dead Cell Removal Kit (Miltenyi Biotec, cat. no. 130-090-101) according to the manufacturer’s instructions with slight modifications described as follows.

#### Peripheral blood mononuclear cells (PBMCs)

PBMC samples characteristically possessed adequate initial cell concentrations and required MACS^TM^ separators for smaller cell samples. Each sample was thus processed for dead cell removal using an MS column (Miltenyi Biotec, cat. no. 130-042-201) and a MiniMACS^TM^ separator (Miltenyi Biotec, cat. no. 130-042-102) on a MACS^TM^ MultiStand (Miltenyi Biotec, cat. no. 130-042-303). Following the initial cell count, samples requiring dead cell removal were washed a second time at 300 *xg* for 10 minutes at room temperature. The supernatant was discarded and the cell pellet was resuspended in 100 µl of Dead Cell Removal MicroBeads per 10^7^ cells, and incubated for 15 minutes at room temperature to magnetically label dead and early apoptotic cells. During the incubation step, 20x Binding Buffer supplied in the dead cell removal kit was diluted with molecular grade water to make a 50 mL 1x stock solution which was then degassed to remove any entrapped air that could result in suboptimal binding to the column. Degassed Binding Buffer (500 uL) was added to a 70 µm pre-separation filter (Miltenyi Biotec, cat. no. 130-095-823) fitted atop a MS Column to primer and pre-moisten the filter. Upon completed incubation, an additional 400 µl of Binding Buffer was added to the labeled cell suspension and mixed well. A 15mL conical tube was placed below the column and 500 µl of cell suspension applied to the moistened filter to limit passage of debris and cell clumps that would otherwise clog the column. The flow-through containing the unlabeled live cells was collected, and four column washes with 500 µl of Binding Buffer were subsequently recovered and combined for a final effluent volume of 2.5 mL.

#### Lymph node

Liquid lymph node (LN) biopsies were similarly processed as described previously for PBMCs, with subtle differences attributed to cellular concentration and dramatically reduced viability. To account for a greater percentage of dead cells and lower initial cell concentrations, a minimum of four vials of LN liquid biopsies were combined for processing. The following details changes impacted by the aforementioned observations and difference in sample type.

A minimum of three LN vials per sample time point were thawed in a 37°C water bath and each carefully passed through a moistened 70 µm pre-separation filter into a 15 mL conical tube. Each vial was rinsed with 1 mL of DPBS to recover any residual cells, and subsequently combined for a total volume of 6 mL. The sample was centrifuged at 400 *xg* for 5 minutes at 4°C. The supernatant was discarded and cell pellet resuspended in 1 mL of DPBS which was then transferred to a 1.5 mL microfuge tube. The cell suspension was washed in the 1.5 mL tube by centrifuging at 300 *xg* for 10 minutes at 25°C. The supernatant was discarded and cell pellet resuspended in 100 µl of Dead Cell Removal MicroBeads as previously stated, and incubated for 15 minutes at room temperature. To accommodate the higher volume of cells going into dead cell removal, an LS column (Miltenyi Biotec, cat. no. 130-042-401) was used with a MidiMACS^TM^ separator (Miltenyi Biotec, cat. no. 130-042-302). Binding Buffer solution and a pre-separation filter were prepared as previously described for PBMCs. The LS column was primed by rinsing with 3 mL of Binding Buffer. Upon completed incubation, an additional 400 µl of 1x Binding Buffer was added to the labeled cell suspension and mixed well. A 15mL conical tube was placed below the column and 500 µl of cell suspension passed through the filter and LS column. The flow-through containing the unlabeled live cells was collected and four column washes with 3 mL of Binding Buffer were collected and combined for a final flow-through volume of 12.5 mL.

Following completion of dead cell removal, cell counts were performed on both PBMC and LN collected effluents to determine total number of cells and ensure greater than 80% viability. The effluents of both sample types were mixed well with a wide bore pipette tip and their volumes alternatively used as dilution factors for cell counting. Cell counts as well as all steps thereafter followed the methods described above in single cell suspension preparation.

### Single-cell DNA sequencing (scDNA-seq)

*Single-cell microfluidics*. The Tapestri® single-cell DNA sequencing V2 platform developed by Mission Bio was performed as per manufacturer’s instructions. Briefly, single-cell suspensions as processed above, were loaded into the Tapestri® instrument and cells individually encapsulated via oil emulsion with a protease enzyme mix. The cells were then lysed and digested in a thermal cycler (Bio-Rad) to enable access to DNA. The encapsulated droplets containing histone-free DNA were then merged with a second droplet comprised of barcoding beads, PCR reagents and primers specific to the custom SIV amplicon panel (Table S2). Regions of interest defined by amplicon design, were subjected to target amplification in a thermal cycler and simultaneously tagged with a unique cell barcode according to the following program: 6 min at 98°C; 10 cycles (30 s at 95°C, 10 s at 72°C, 3 min at 61°C, 20 s at 72°C); 12 cycles (30 s at 95°C, 10 s at 72 °C, 3 min at 48°C, 20 s at 72°C); 2 min at 72°C. Emulsions were broken and samples stored at -20°C for less than six months before advancing to cleanup of PCR products.

Library cleanup proceeded with PCR product digestion using a thermocycler, followed by Ampure XP bead-based cleanup. In short, a 0.72X Ampure XP reagent (Beckman Coulter, cat. no. A63881) ratio was initially incubated with the digested PCR products. The solution was placed on a magnet and beads allowed to separate. The beads containing adhered DNA were washed twice with freshly made 80% ethanol then allowed to air dry until all residual ethanol was removed. The beads were then resuspended in nuclease-free water and further purified on the magnet. The High Sensitivity dsDNA 1X kit and Fluorometer (Qubit^TM^ 4 Fluorometer, Invitrogen) was used to quantify the purified PCR product. Yields of DNA between the recommended 0.2 ng/µl to 4.0 ng/µl range were confirmed. A targeted library PCR step was then performed to add unique dual index sequences to the amplicons. Distinct pairings of index primers were used and recorded for separate samples to allow for multiplex sequencing. Upon completion of targeted library PCR, a second library cleanup was completed using a 0.69X Ampure XP bead ratio, and all other cleanup steps similarly executed as in the previous library cleanup. The final DNA library concentration was again quantified by fluorometry and purity of on-target library analyzed with the Agilent 4150 TapeStation system (cat. no. G2992AA). For samples observed with >30% of smaller off-target product, further library purification using gel-size selection was carried out prior to sequencing. Constructed libraries were pooled and sequenced using various Illumina platforms. Sequencing was performed with an Illumina NovaSeq 6000 SP Reagent kit v1.5 in 2 x 250 bp paired-end mode, on the NextSeq 1000/2000 System with a NextSeq 1000/2000 P1 reagent kit generating 2 × 250 bp paired-end reads, or on the MiSeq system for 500 cycles. The Tapestri® instrument protocol version referenced in this study was PN_3354H.

#### Gel library purification

. Size selection of the library DNA was performed using an 11 cm by 14 cm 2% agarose gel with 8 µl of Syber safe gel dye. The samples for multiplexed sequencing that contained >30% off-target small fragments were pooled using equimolar quantities of on-target library prior to size selection. The gel was loaded with 25 µl of the pooled sample and 5 µl of 6x orange DNA loading dye-Fermentas into a single well. A volume of 15 µl of GeneRuler 1kb DNA ladder (ThermoFisher Scientific) was used for size identification of the band size of interest. The gel was then run at 64 V for 2 hours. The band corresponding to the size of interest at ∼450bp was cut and purification was performed using the QIAquick Gel Extraction Kit (Qiagen) as per the manufacturer’s instructions.

### Bioinformatic pipeline optimization

The commercially available Tapestri bioinformatic pipeline was used with the following modifications: uniformity and completeness thresholds were made to account for expected differences in largely “working amplicons” for human DNA, in which the pipeline was built to handle, as opposed to the breadth of variation experienced for viral DNA due to cumulative mutations and deletions. Animal-specific sequence alignments, scripts, as well as specifics on how and what pipeline parameters were changed, can be found at https://github.com/salemilab/scDNA-seq.

### Near full-length viral genome assembly

Cell-specific barcodes containing reads mapping to the custom SIV reference genome were extracted and split from the rest of the cells, and individual bam files created for each barcode using SAMTOOLS V1.12^64^. Consensus reconstruction using SAMTOOLS V1.12 and ivar^65^ v1.3.1 resulted in FASTA files for individual genomes that were then concatenated and aligned with VIRALMSA^66^ to the in-house reference sequence for downstream analysis. Cells were considered SIV+ based on a criteria of possessing a minimum depth of 10 reads for at least two consecutive amplicons. All batch scripts to obtain the final consensus sequences can be found at https://github.com/salemilab/scDNA-seq.

### Construction of custom plasmids

Two plasmids were custom-designed by our group and synthesized directly by GenScript. Construction of the plasmids included a pUC18 backbone with ampicillin resistance and either the wild-type (WT) in-house reference genome or a hypermutated version with 1,481 hypermutations, comprising all possible GG|GA dinucleotide sets.

### Plasmid transfection of 293T cells

293T cells were obtained from BEI and maintained at 37°and 5% carbon dioxide in high glucose DMEM (cat. no. 10-013-CV) with 10% heat-inactivated FBS, 10,000 µg/mL, and penicillin 10,000 U/mL. 293T cells were seeded at 4.0 x 10^5^ cells per well in a 12 well plate. When cells reached 80-90% confluence (around 24 hours of seeding), cells were transfected with 250 ng of a single plasmid per well using Lipofectamine 2000 (invitrogen cat. no. 11668-030) in a ratio of 1:1 Lipofectamine:DNA and 250 µl Opti-MEM (invitrogen cat. no. 31985062) per well, added in a slow drop-wise manner. Media was replaced (with fresh) 1 hour before transfection and 24 hours post-transfection. Cells were harvested at 24 hours post-transfection.

### Plasmid extraction from HEK293T cells

For isolation of DNA from the transfected cells, the DNeasy Blood & Tissue kit (Qiagen cat. no. 69506) was used, according to the manufacturer’s instructions. DNA concentrations were determined fluorometrically using the Qubit^TM^ 4 Fluorometer (Invitrogen) and Qubit^TM^ dsDNA HS Assay kit (Invitrogen).

### Copy number determination via droplet digital PCR (ddPCR)

Droplet digital PCR (ddPCR) was used to corroborate our sequencing measurements. Eleven ddPCR reactions were performed in triplicate including macaque samples (PBMCs and LN at necropsy from monkeys JA41, JP70, and HG02), WT plasmid alone, WT-transfected HEK293T cells, hypermutant plasmid alone, hypermutant-transfected HEK293T cells, and non-transfected cells. The ddPCR Multiplex Supermix (Bio-rad) was used to set up all 6-plex reactions following manufacturer’s instructions. The following primers and probes were used: *Psi*_*F* : 5*′ −CGGAGTGCTCCTATAAAGGC−*3*′, Psi*_*R* : 5*′ −CAAGACGGAGTTTCTCGC−*3*′, Psi_p_robe* : 5*′ −FAM− CCGGTTGCAGGTAAGTGCAACA −* 3*′, HIV* _*Env*_*F* : 5*′ − GTGGCACCTCAAGAAATAAAAGAG −* 3*′, HIV* _*Env*_*R* : 5*′ − AATAAAGTCCGGGACTGAGC −* 3*′, HIV* _*Env*_*probe* : 5*′ −HEX −TTTTCTCGCAACGGCAGGTTCT −* 3*′, HIV* _*Pol*_*F* : 5*′ −GAGAGAAAGCAGAGAGAAGCC −* 3*′, HIV* _*Pol*_*R* : 5*′ −AGGCTGTCCTTCAATATGAGC −* 3*′, HIV* _*Pol*_*probe* : 5*′ − ATTO*590 *−AGGTGACAGAGGATTTGCTGCA−* 3*′, SIV* _*Pol*_*F* : 5*′ −CAGGGATAGAGCACACCTTTG−* 3*′, SIV* _*Env*_*F* : 5*′ −CCTCAATAAAGCCTTGTGTAAAATTATC−*3*′, SIV* _*Env*_*R* : 5*′ −GCTGTTGTTGTTGATGATTTTGTC−*3*′, SIV* _*Env*_*probe* : 5*′ −Cy*5.5*−CCCCATTATGCATTACTATGAGATGC−*3*′, SIV* _*Pol*_*R* : 5*′ −GCAATGAACTGCCATTAATACTATGG−*3*′, SIV* _*Pol*_*probe* 5*′ −Cy*5 *−TACCATACAATCCACAGAGTCAGGAG−* 3*′, RPP*30_*F* : 5*′ −GATGCTCCGGGAGTATGTAAC −* 3*′, RPP*30_*R* : 5*′ −CTCCCTGCTTGTCACCTATATAAC −* 3*′, RPP*30_*probe* : 5*′ −ROX −CAAGCTGGGAGACGGAAGAGTCAG−* 3′. Re-striction enzyme NheI-HF (NEB) was used to linearize plasmids. After generating droplets on the QX600 droplet generator (Biorad), reactions were thermal cycled at 95 °C for 10 min, followed by 40 cycles of 94 °C for 30 s and 59 °C for 2 min, with a final single step at 98 °C for 10 min. Samples were held at 4 °C after the reaction was completed and before droplets were read. Droplets were read with the QX600 Droplet Reader (Biorad) and droplet counts were determined using the QX Manager Standard Edition software (Biorad).

### Classification of viral genomes

Viral DNA genome sequences were categorized as “deletion-defective”, “hypermutation defective,” or intact. To combat erroneous classifications as deletion defective owing to potential variability in primer-binding sites and therefore loss of sequenced reads, absence of four consecutive amplicons were used. Deletion-defective virus were further sub-classified by having either 3’ only deletions, 5’ only deletions, or double deletions comprising both ends. For these three sub-categories, the first two amplicons within the 5’ and/or 3’ regions were required to be missing. Hypermutated genomes were classified using Hypermut2.0^67^.

### Genetic diversity calculations

Hypermutated sequences were identified using Hypermut2.0 (available from the Los Alamos National Laboratory HIV sequence database (https://www.hiv.lanl.gov) and hypermutated sites masked through replacement with ’N’, similar to Shahid *et al.*, 2024^48^. Calculations for population genetic diversity were performed in R^68^ using the APE V5.6-2 package^69^ function dist.dna to compute matrices of pairwise distances using the TN93 model of nucleotide substitution. *TRIM*5*α* diversity was assessed using mutational frequencies at base positions of the targeted macaque *TRIM*5*α* gene (chromosome 14, exon 8).

### Phylogenetic analyses

Likelihood mapping and maximum likelihood tree reconstruction were performed using IQ-TREE V2.1.3 according to the best-fitting nucleotide substitution model. Following 100 traditional (Felsenstein^70^) bootstrap replicates, estimation of the transfer bootstrap expectation (TBE) was performed on bootstrapped trees using the BOOSTER module, as described previously^49^). Robinson-Foulds distances^50^ were calculated and normalized in R^68^ using the PHANGORN V2.8.1 package^71^ RF.dist function in TREEDIST. Joint likelihood reconstruction of ancestral tissue states was performed using the ACE function with ER model in the R APE package. Resulting tree files are available in https://github.com/salemilab/scDNA-seq.

## Data and Code Availability

Raw SIV sequence data that support the findings of this study have been deposited in GenBank under the accession code XXXXXXXX. Scripts concerning viral genome and *TRIM*5*α* consensus reconstruction are available on the scDNA-seq GitHub along with scripts for generating the phylogenetic trees presented in the manuscript main text (https://github.com/salemilab/scDNA-seq). Guidance on setting pipeline parameters are additionally provided, as well as all recovered SIV sequences obtained in this study in FASTA format.

## Acknowledgements

Research reported in this manuscript was supported by the National Institutes of Health (NIH) National Institute of Neurological Disease and Stroke [R01NS063897 to M.S.] and the Stephany W. Holloway University Chair in AIDS Research. Sequencing was provided by the University of Louisville Sequencing Technology Center, supported in part by the National Institute of General Medical Sciences [P20GM103436 (KY INBRE)].The authors also acknowledge the University of Florida and University of Louisville Research Computing for providing computational resources and support that have contributed to the research results reported in this publication.

## Author information

### Contributions

L.D., A.R.M., M.C., J.E., A.B., and F.R. contributed to the experimental aspects of the study, S.D.S. and B.R.M. to the development of analytical tools and analysis of data, and S.L.K.P., M.S., and B.R.M. to the supervision of experimental and analytical components. B.R.M. conceived of the presented idea and verified experimental and analytical methods. All authors discussed the results and contributed to the final manuscript.

### Corresponding authors

Correspondence to Brittany Rife Magalis or Marco Salemi

## Supplementary

**Table S1.** PCR conditions for viral load quantification.

| Step | Temp (°C) | Time (mm:ss) | Number of cycles |
| --- | --- | --- | --- |
| Initial Denaturation | 95 | 10:00 | 1 |
| Denaturation | 94 | 00:15 | 40 |
| Polymerase | 62 | 01:00 | 40 |

**Table S2.**
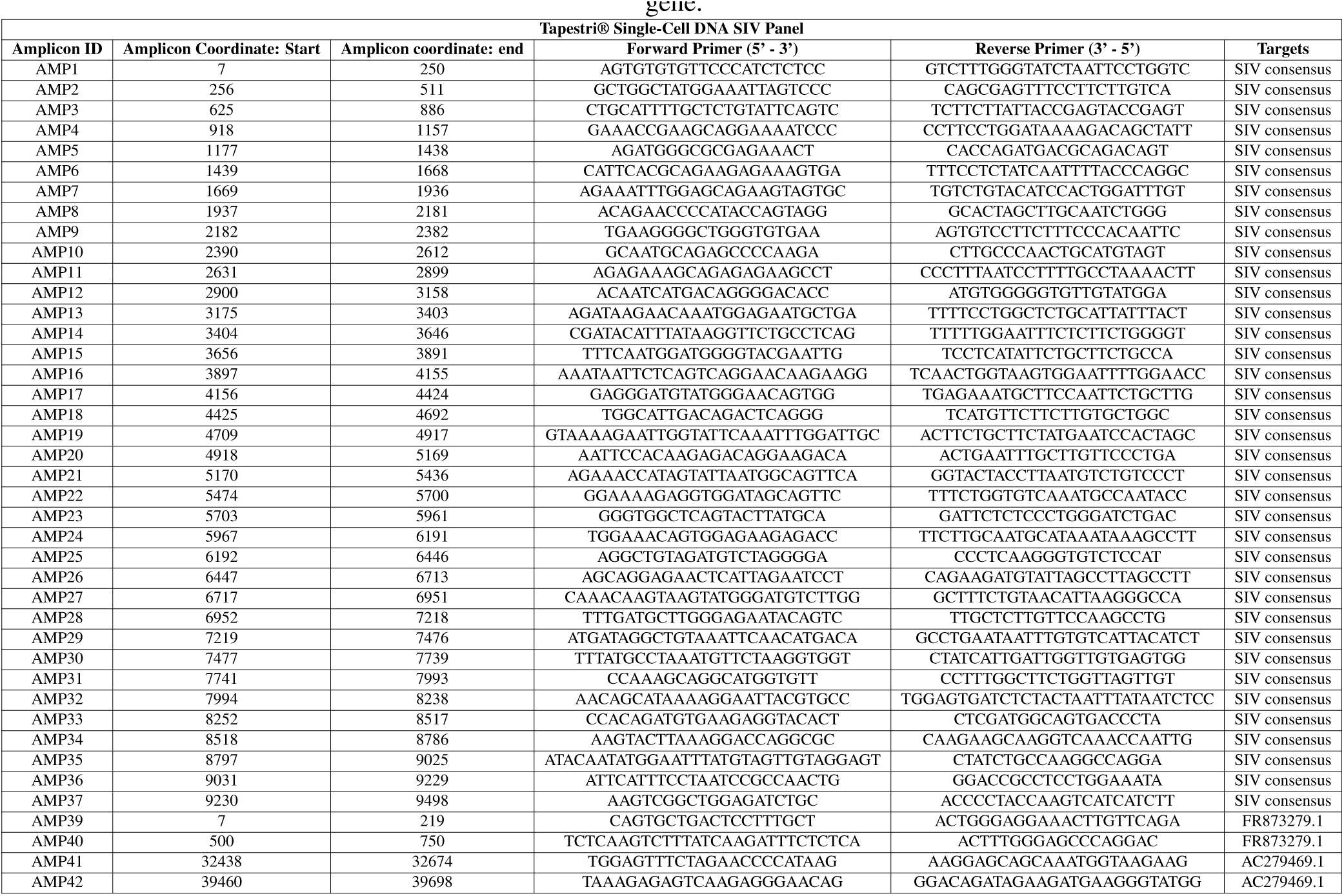
Primers used in scDNA-seq of SIV genome and Rhesus macaque *RPP*30 and *TRIM*5*α* genes. Amplicon IDs are listed with their forward and reverse primer sequences and corresponding locations within the reference genome. Amplicons 1 − 38 correspond to regions within the custom reference sequence and U5U3 junction. Amplicons 39 and 40 correspond to macaque *RPP*30, and Amplicons 41 and 42 to the SPRY domain of the macaque *TRIM*5*α* gene.

**Table S3.**
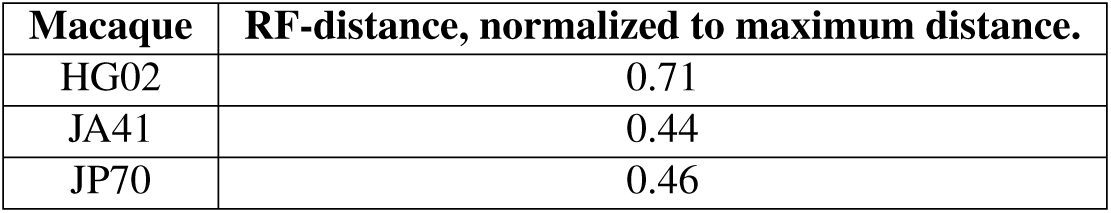
Degree of similarity in topological symmetry between full genome and *env gp120* phylogenies for each of three macaques. Genome sequences containing full-length *gp120* for each animal were used in the reconstruction of a maximum likelihood phylogeny for the full sequence and *gp120* segment alone. Robinson-Foulds distances (RF-distance) were calculated, describing the degree of similarity between tree topologies, with 05 indicating identical topologies and 100% representing random taxon assignment.

| Macaque | RF-distance, normalized to maximum distance. |
| --- | --- |
| HG02 | 0.71 |
| JA41 | 0.44 |
| JP70 | 0.46 |

**Figure S1.**
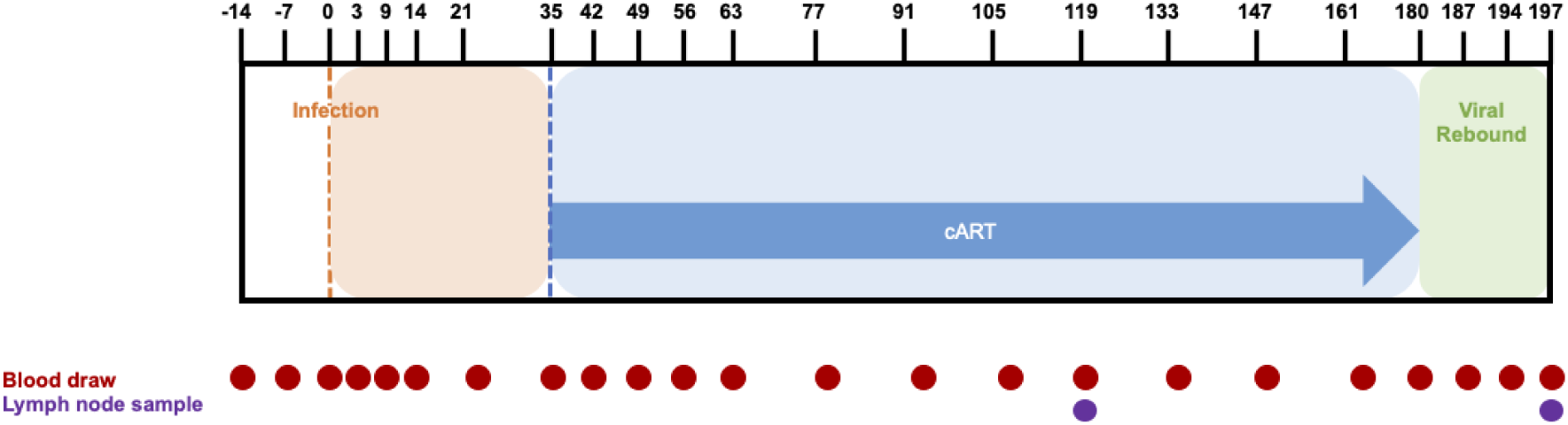
Timeline of events for the macaque model of HIV infection used in this study. *Rhesus* macaques were infected with a pathogenic strain of SIV (SIVmac251), represented throughout this study as day 0. Blood and lymph node samples were taken at pre-specified time points during cART and following cART interruption for viral load determination and/or single-cell sequencing.

**Figure S2.**
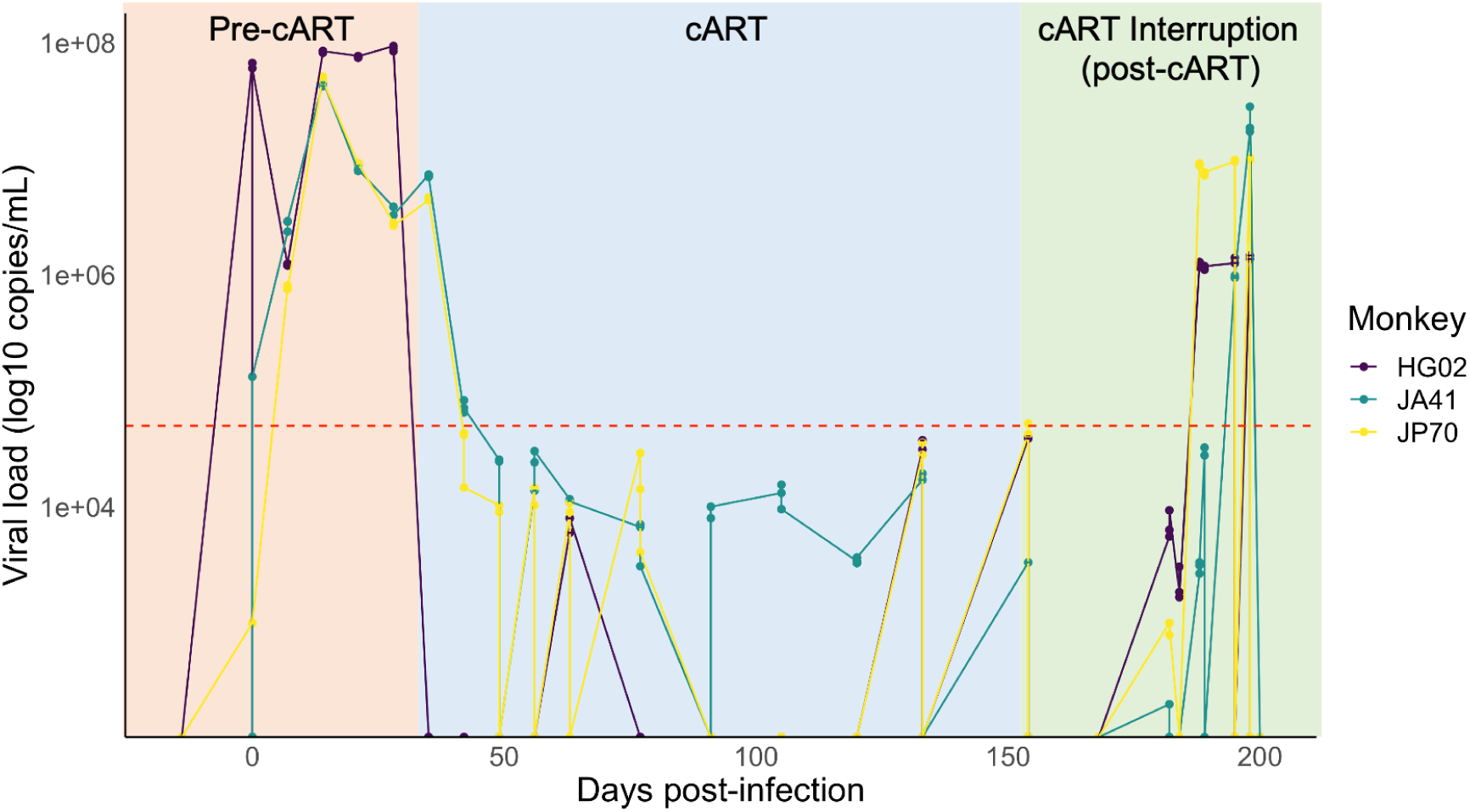
Viral load (copies/uL) over time (days post-inoculation) in three macaques. Quantitative polymerase chain reaction was used to measure viral load, with a reliable detection limit (red dashed line, 50 copies/uL) determined using a standard curve.

**Figure S3.**
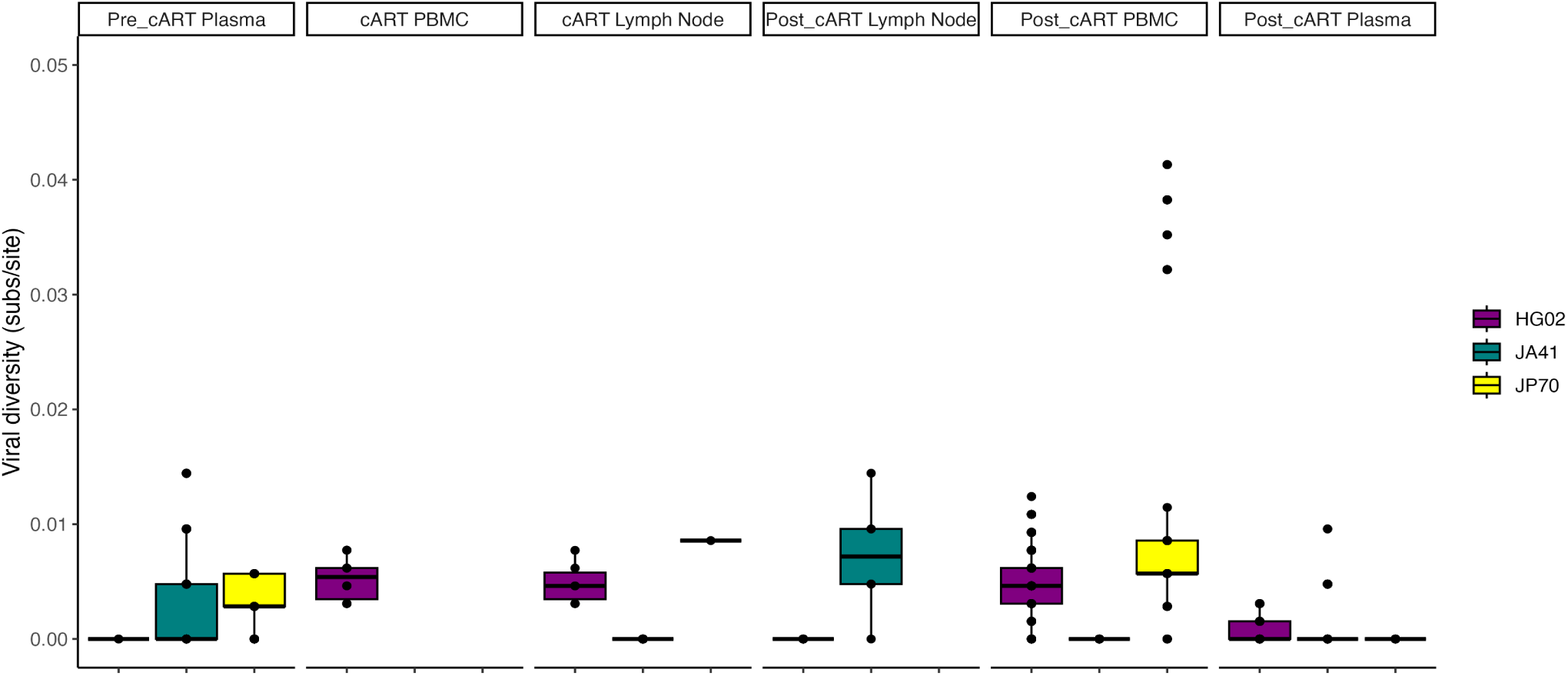
Pairwise genetic diversity across SIV DNA genomes. Pairwise genetic distances among SIV DNA sequences for each animal, tissue, and time point was estimated using the TN93 nucleotide substitution model. Data median and 25th and 75th percentiles are represented, and colors represent individual animals (legend at right).

**Figure S4.**
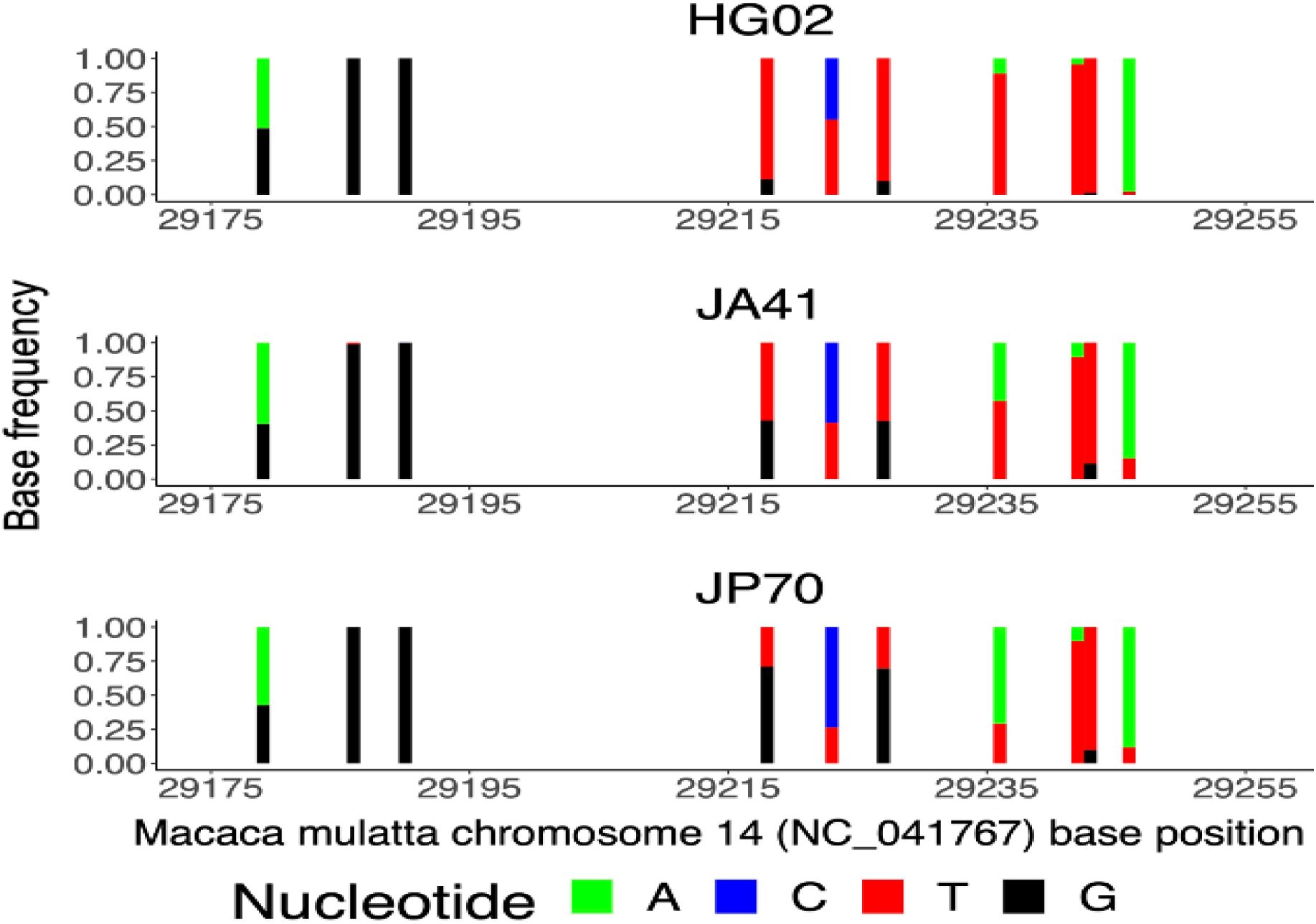
*TRIM*5*α* genetic diversity. Multi-allelic site frequencies are reported among *TRIM*5*α* sequences for each of three macaques. Colors represent individual bases (legend at bottom).

**Figure S5.**
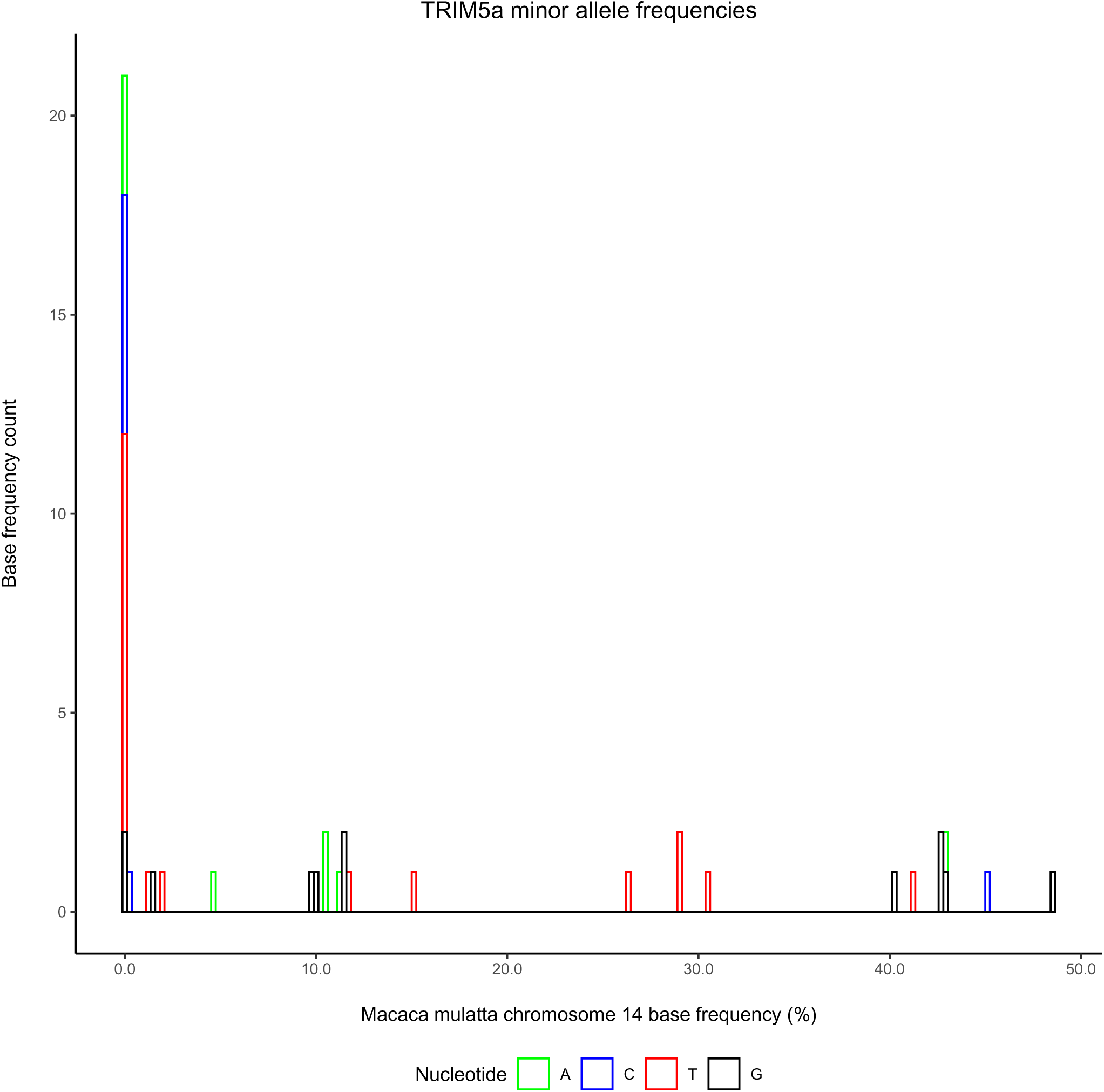
Distribution of TRIM5a minor allele frequencies. Minor allele nucleotide bases are distributed by percent frequency observed in amplicon 42 of the *Rhesus* macaque TRIM5a chromosome 14. Base counts are stacked to showcase nucleotide prevalence at all indicated frequencies.

**Figure S6.**
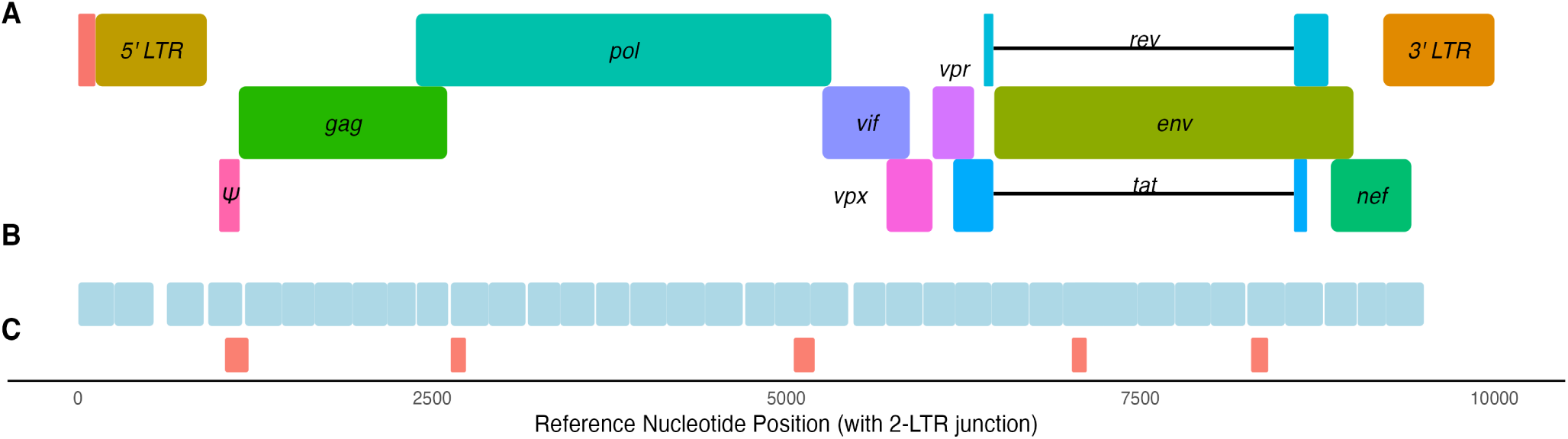
SIV genome coverage with scDNA-seq and ddPCR. **(A)** SIV genome map with coordinates (x-axis) representing the reference sequence including 2-LTR junction at nucleotide sites 7-250. **(B)** scDNA-seq amplicon coverage across SIV reference genome. **(C)** ddPCR amplicon coverage cross the SIV reference genome, designed to be complementary to scDNA-seq amplicons

**Figure S7.**
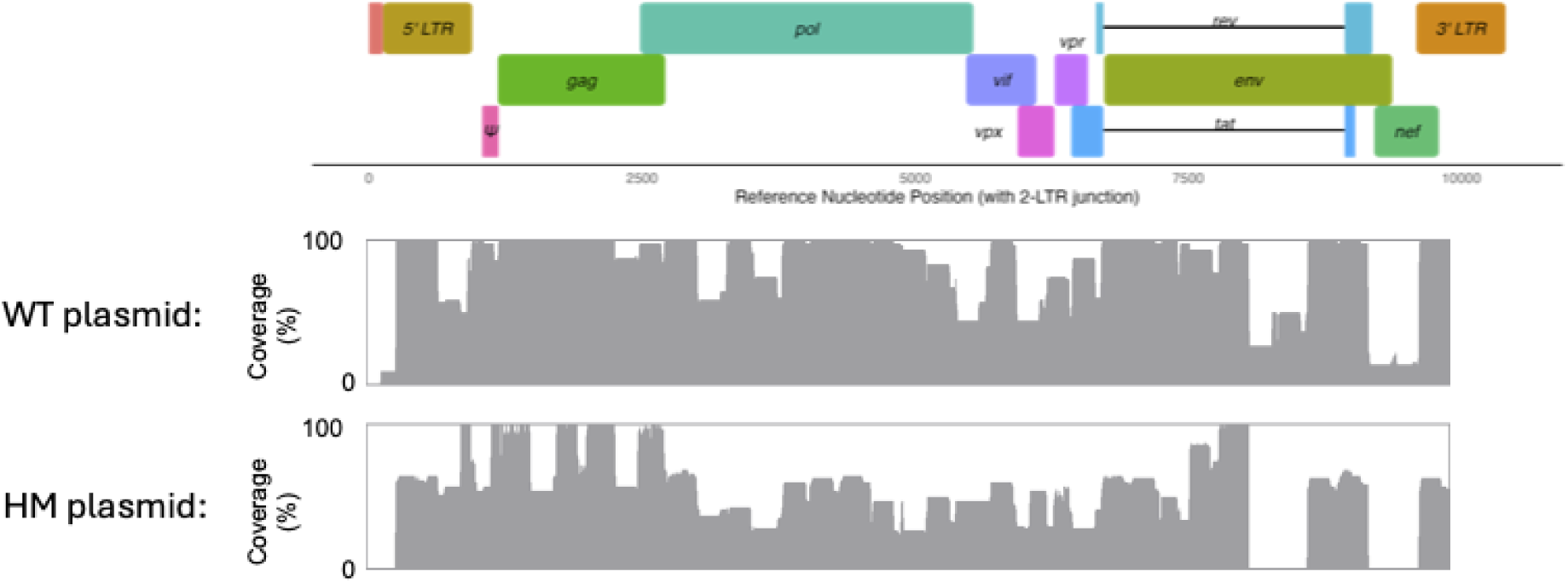
Amplicon coverage across wild type (WT) and hypermutant (HM) SIV plasmid genomes transfected into HEK293T cells.

**Figure S8.**
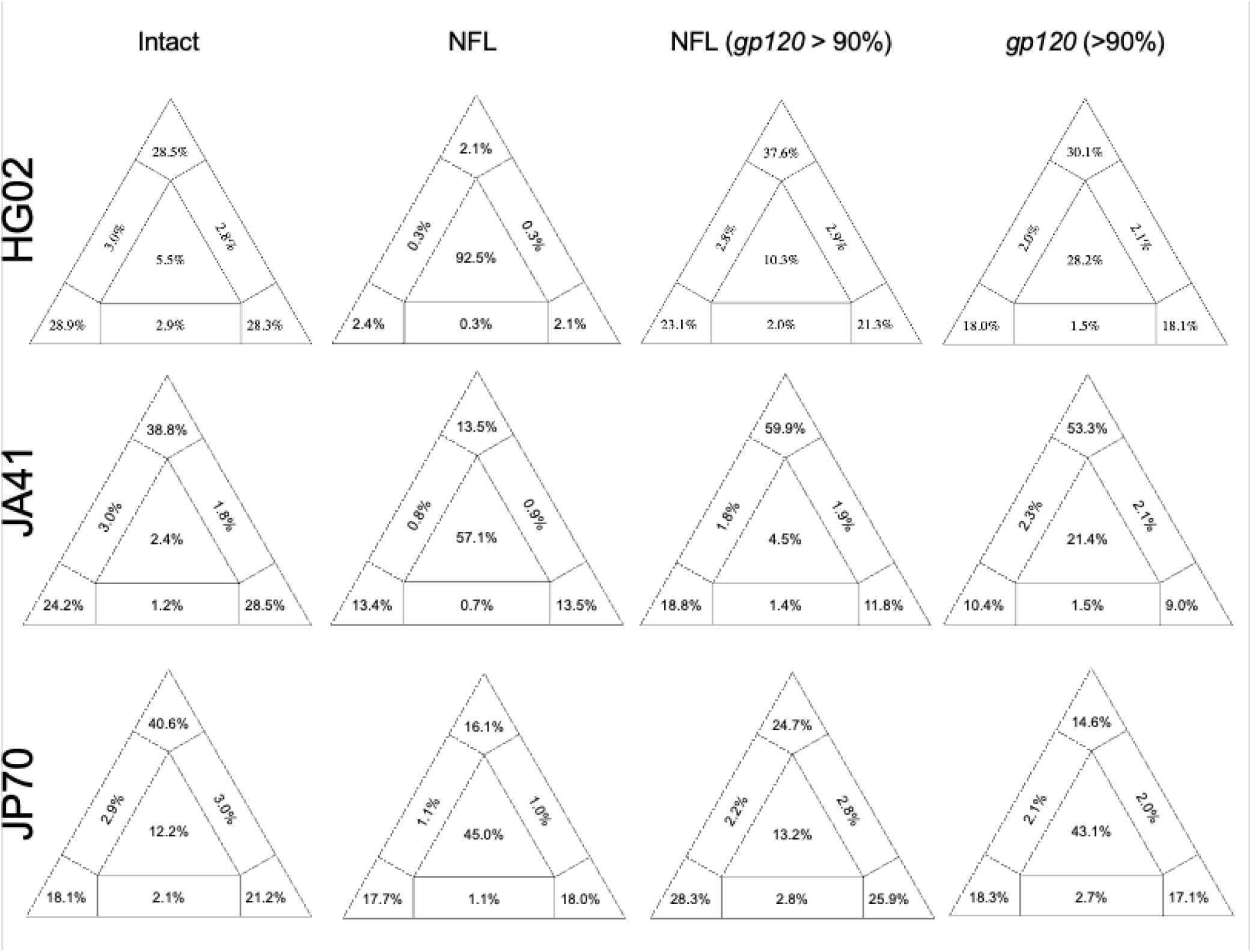
Comparison of phylogenetic resolution for varying sequence inclusion scenarios (columns) within each animal (row). Phylogenetic likelihood maps demonstrate the percentage of sampled sequence quartets classified as phylogenetically resolved (corners), partially resolved (outer edges), or unresolved (center) during phylogenetic reconstruction.

**Figure S9.**
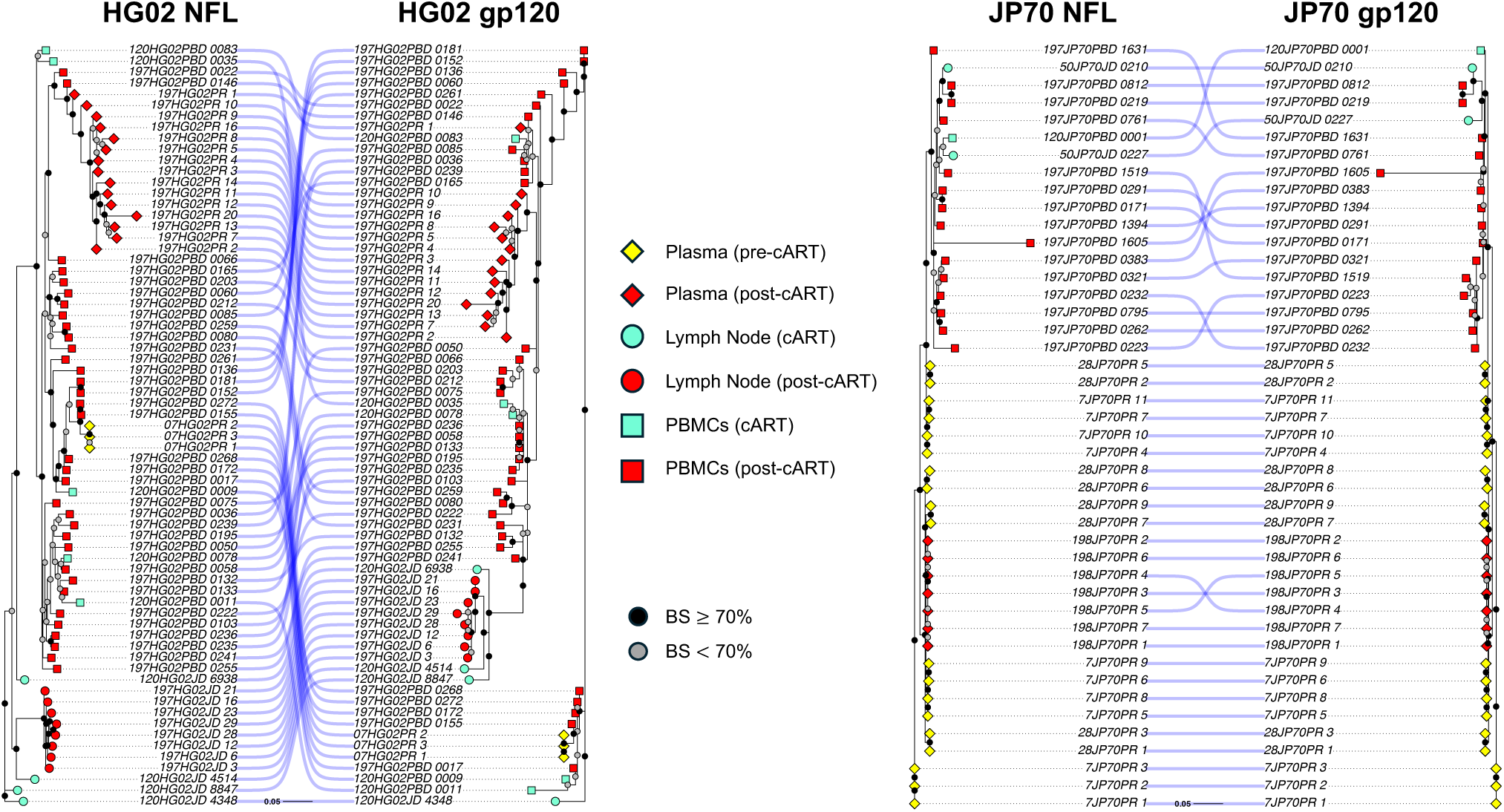
Comparison of phylogenetic placement for near-full genomes and subgenomic *env gp120* for animals HG02 (left) and JP70 (right). Maximum likelihood phylogenetic trees were reconstructed using IQ-TREE V2^72^ using the best-fitting nucleotide substitution model. Shape indicates tissue type (square: PBMC, circle: LN), while color indicates time point (cyan: cART, red: post-cART). Support for branching events was provided through bootstrapping^49^, with *≥* 70% considered well-supported and represented by a black circle and < 70% represented by a gray circle.

